# Sleep-dependent engram reactivation during hippocampal memory consolidation is associated with subregion-specific biosynthetic changes

**DOI:** 10.1101/2022.11.09.515837

**Authors:** Lijing Wang, Lauren Park, Weisheng Wu, Dana King, Alexis Vega Medina, Frank Raven, Jessy Martinez, Amy Ensing, Katherine McDonald, Zhongying Yang, Sha Jiang, Sara J. Aton

## Abstract

Post-learning sleep is essential for hippocampal memory processing, including contextual fear memory (CFM) consolidation. We used targeted recombination in activated populations (TRAP) to label context-encoding engram neurons in the hippocampal dentate gyrus (DG) and assessed reactivation of these neurons after fear learning. Post-learning sleep deprivation (SD), which impairs CFM consolidation, selectively disrupted reactivation of inferior blade DG engram neurons. This change was linked to more general suppression of neuronal activity markers in the inferior, but not superior, DG blade by SD. To further characterize how learning and subsequent sleep or SD affect these (and other) hippocampal subregions, we used subregion-specific spatial profiling of transcripts and proteins. We found that transcriptomic responses to sleep loss differed greatly between hippocampal regions CA1, CA3, and DG inferior blade, superior blade, and hilus – with activity-driven transcripts, and those associated with cytoskeletal remodeling, selectively suppressed in the inferior blade. Critically, learning-driven transcriptomic changes, measured 6 h following contextual fear learning, were limited to the two DG blades, differed dramatically between the blades, and were absent from all other regions. These changes suggested an increase in glutamatergic receptor signaling, and a decrease in GABA receptor signaling, during CFM consolidation. Protein abundance across hippocampal subregions was also differentially impacted by learning and sleep, with most alterations to protein expression restricted to DG. Together, these data suggest that the DG is critical for sleep-dependent memory consolidation, and that the effects of sleep loss on the hippocampus are highly subregion-specific, even within the DG itself.

## Introduction

Hippocampal memory consolidation is affected by the amount and quality of post-learning sleep (1–4). In both animal models and human subjects, sleep deprivation (SD) negatively impacts consolidation of hippocampus-dependent memories (5–7). For example, in mice, consolidation of contextual fear memory (CFM), a canonical form of Pavlovian conditioning (8) is disrupted by SD in the first 5-6 h following single-trial contextual fear conditioning (CFC; pairing exploration of a novel context with a foot shock) (9–13). Recent studies have shown that SD can alter basic features of hippocampal network function, including oscillatory patterning of network activity (11), intracellular signaling (10, 14, 15), transcription and translation (12, 16–19), and excitatory-inhibitory balance (20, 21). However, the precise mechanisms responsible for SD-driven disruption of hippocampal memory storage remain unknown.

Growing evidence suggests that CFM is encoded via activation of a sparse population of hippocampal engram neurons (22). Natural cue-driven memory recall (i.e., upon return to the conditioning context) can reactivate at least some of the same neurons active during CFC in hippocampal structures including DG (23, 24). Optogenetic reactivation of these so-called engram neurons drives fear memory retrieval (25, 26). Based on these findings, offline reactivation of engram neurons is widely hypothesized to serve as the mechanistic basis of memory trace storage. Indeed, recent data suggest that consolidation is associated with, and requires, sleep-associated reactivation of engram populations in neocortex (27, 28) and hippocampal area CA1 (29). However, we have recently found that SD profoundly disrupts network activity in DG (12, 17, 20, 21). This effect is mediated in part through acetylcholine-dependent activation of somatostatin-expressing interneurons in DG - which in turn suppress activity among DG granule cells (20). DG is a critical input structure to the hippocampus (receiving input from neocortex via the entorhinal cortex), and DG engram neurons’ activity is necessary and sufficient for CFM recall (25, 26, 30). Thus, a critical unanswered question is how post-CFC sleep and SD affect post-learning engram neuron reactivation in the context of CFM consolidation.

We first addressed this question using targeted recombination in activated populations (TRAP) (31) to label CFC-activated engram neurons in dorsal hippocampal DG. We find that these TRAP-labeled engram neurons selectively reactivate over the first few hours following CFC. Post-learning SD disrupts this offline reactivation in a region-specific manner, preventing reactivation specifically in inferior blade DG granule cells. These findings suggest a subregion-specific, instructive, sleep-dependent mechanism for CFM consolidation, which selectively drives engram neuron reactivation in one subregion (inferior blade of DG). To identify subregion-specific cellular mechanisms associated with this phenomenon, we used spatial transcriptomics to identify transcript changes associated with CFC and subsequent sleep or SD in subregions of DG, CA1, and CA3. Surprisingly, SD-driven transcriptomic changes differed substantially between these subregions and varied dramatically between the two DG blades. Many of the changes occurring in the inferior blade with SD could play a causal role in preventing engram neuron reactivation during SD, or the functional consequence of engram neuron activity disruption – including changes to pathways involved in regulating synaptic structure. Moreover, learning-associated transcriptomic changes: 1) were present only in the two DG blades 6 h following CFC, 2) were absent from all other subregions profiled at this timepoint, and 3) differed significantly between the blades. Further characterization of hippocampal subregions with spatial protein and phosphoprotein profiling again showed distinct, subregion-specific effects of both learning and subsequent sleep vs SD. Together, these findings reveal previously uncharacterized heterogeneity and subregion-specificity in the effects of both learning and sleep on the hippocampus. The present data provide new insights into mechanisms by which post-learning sleep contributes to hippocampal memory consolidation.

## Results

### Hippocampal engram neuron reactivation during post-learning sleep is subregion- specific

To visualize neurons activated in the hippocampus by a learning experience, we used a genetic strategy recently used to identify visual memory engram neurons in mouse primary visual cortex (V1) (27). To identify context-activated neurons with TRAP (31), *cfos-CRE^ER^* transgenic mice were crossed to a *tdTomato* reporter line (*cfos::tdTomato*). At lights on (i.e, the beginning of the rest phase; Zeitgeber time [ZT]0), *cfos::tdTomato* mice were placed in a novel context (Context A) for 15 min of free exploration, immediately after which they were administered 4-hydroxytamoxifen (4-OHT) to label context-activated hippocampal neurons with tdTomato. 6 days later at ZT0, the mice were either returned to Context A (A to A) or placed in a dissimilar Context B (A to B) for 15 min. 90 min after the second period of exploration (**Fig. 1a**), mice were perfused and hippocampal cFos expression was quantified to assess neuronal activity. In agreement with previous reports using a different transgenic strategy (24)(26), TRAPed tdTomato+ neuronal cell bodies in the DG granule cell layers were more likely to be reactivated during Context A re-exposure than during exploration of Context B (**Fig. 1b-c**). Conversely, a larger proportion of A to A cFos+ neurons were tdTomato+ compared with A to B cFos+ neurons (**Fig. 1e**). We next compared the specificity of Context A engram neuron activation in the superior vs. inferior blades’ granule cell layer within DG. In the superior blade, but not the inferior blade, the proportion of cFos+ tdTomato+ neurons was significantly higher in the A to A group compared with A to B (**Fig. 1d,f**), despite the average proportion of cFos+ tdTomato+ neurons being similar between the two blades for animals in the same experimental condition (e.g., A to A). These differences between the A to A and A to B were not due to either the number of total tdTomato+ neurons or total cFos+ neurons, which were similar between the two groups, both across the entire granule cell population (**Fig. S1a-b**) and within each of the two blades (**Fig. S1c-d**). However, consistent with previous reports (31, 32), tdTomato+ and cFos+ neuron numbers were consistently higher in the superior vs. inferior blade (**Fig. S1c-d**). Together, these data suggest that TRAPed engram cells in DG granule cell layer - particularly in the superior blade - are reliably reactivated upon re-exposure to the same context.

**Fig. 1:**
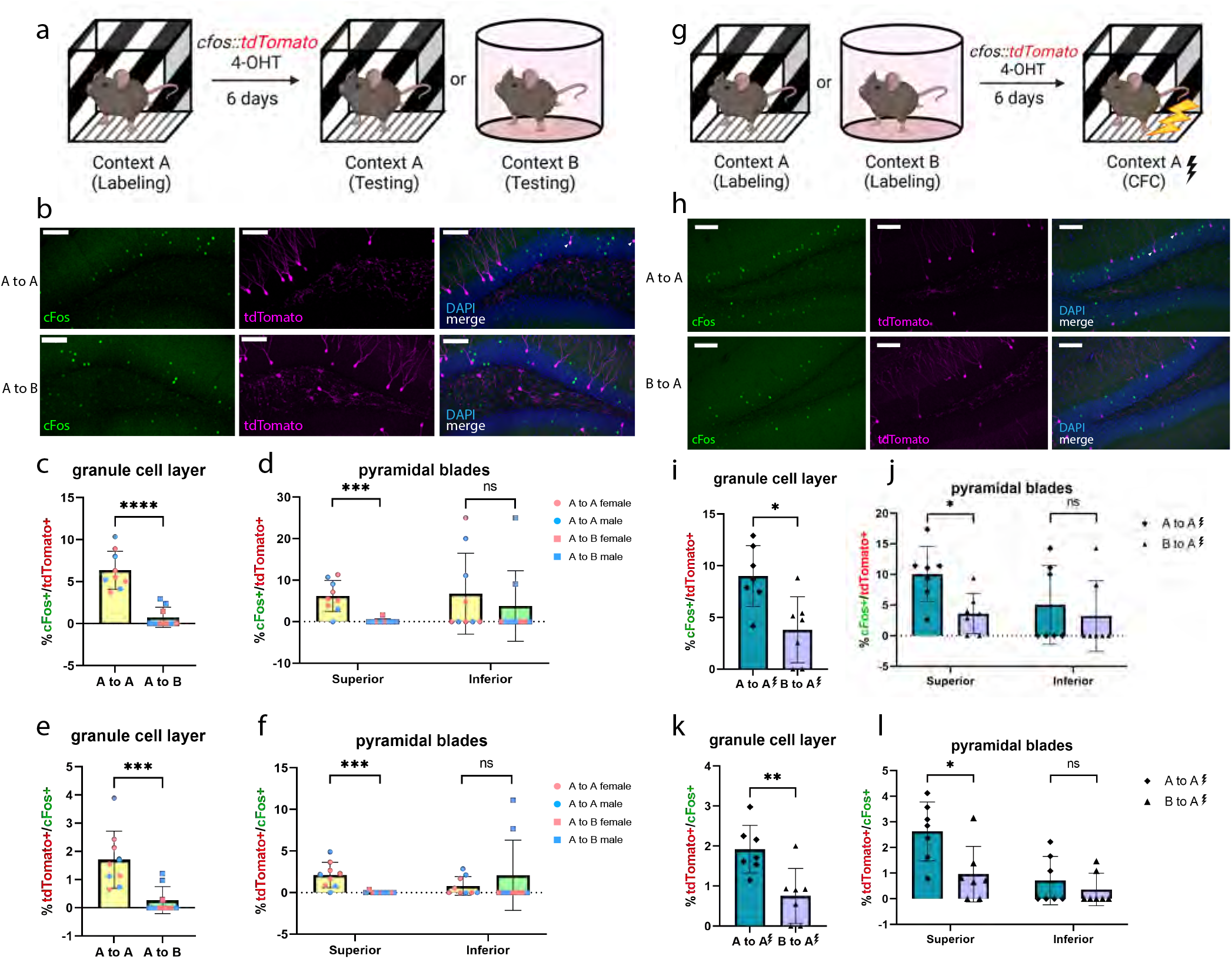
Contextual fear conditioning (CFC) reactivates context-labeled neurons. **a,** Male and female *cfos::tdTomato* mice were injected with 4-hydroxytamoxifen (4-OHT) following Context A exploration. 6 d later, mice were either re-exposed to Context A (A to A) or were placed in a dissimilar Context B (A to B) prior to tissue harvest. **b,** Representative images showing overlap of tdTomato (magenta) and cFos protein (green). Examples of colocalization within a neuron is indicated with a white arrowhead. Scale bar =100 μm. **c,** Percentage of cFos+/tdTomato+ cells in the DG granule cell layer is significantly higher in A to A (*n* = 9; 4 males and 5 females) compared with in A to B (*n* = 9; 5 males and 4 females). **** indicates *p* < 0.0001, Mann Whitney test. **d,** cFos+/tdTomato+ overlap percentage differed between conditions in the superior blade (*** indicates *p* = 0.0006, Mann Whitney test) but not the inferior blade (ns = not significant). **e,** Percentages of tdTomato+/cFos+ cells in the DG granule cell layer. *** indicates *p* = 0.0009, Mann Whitney test. **f,** tdTomato+/cFos+ overlap percentage differed between conditions in the superior (*** indicates *p* = 0.0004, Mann Whitney test) and inferior (ns = not significant) blade of DG granule cell layer. **g,** Male *cfos::tdtomato* mice were injected with 4-OHT following either Context A or Context B exploration. 6 d after labeling, all mice received contextual fear conditioning (CFC) in Context A prior to tissue harvest. **h,** Representative images showing overlap of tdTomato (magenta) and cFos protein (green). Examples of colocalization within a neuron is indicated with a white arrowhead. Scale bar = 100 μm. **i,** Percentages of cFos+/tdTomato+ cells in the DG granule cell layer is significantly higher in A to A (*n* = 7) than B to A (*n* = 7). * indicates *p* = 0.0256, Mann Whitney test. **j,** The cFos+/tdTomato+ overlap percentage is significantly higher in the superior blade (* indicates *p* = 0.0256, Mann Whitney test) but not the inferior blade (ns, not significant). **k,** Percentages of tdTomato+/cFos+ cells in the DG granule cell layer. ** indicates *p* = 0.007, Mann Whitney test. **l,** Percentages of tdTomato+/cFos+ cells in the superior (* indicates *p* = 0.0256, Mann Whitney test) and inferior (ns = not significant) blade of DG granule cell layer. All bars indicate mean ± s.d.

We next tested how reactivation of DG engram populations was affected by context-associated CFC. Male *cfos::tdTomato* mice explored either Context A or dissimilar Context B at ZT0, and were administered 4-OHT to label context-activated neurons. 6 days later at ZT0, mice from both groups underwent single-trial CFC in Context A and were perfused 90 min later to quantify CFC-driven neuronal activation (**Fig. 1g**). Again, a greater proportion of Context A-activated tdTomato+ DG granule cells (vs. Context B-activated tdTomato+ neurons) were cFos+ following CFC in Context A (A to A vs. B to A; **Fig. 1h-i**), and a higher percentage of cFos+ neurons were previously TRAPed tdTomato+ neurons in the A to A paradigm vs. the B to A paradigm (**Fig. 1k**). As was observed previously, in the superior blade, but not the inferior blade, the proportion of cFos+ tdTomato+ neurons was higher following the A to A CFC paradigm compared with B to A CFC paradigm (**Fig. 1 j,l**). Again, the total numbers of tdTomato+ cFos+ neurons were similar between groups (**Fig. S1e-f**), and were more numerous in the superior vs. inferior blades (**Fig. S1g-h**). Thus, CFC selectively reactivates specific context-encoding granule cells in the DG superior blade.

Across experimental groups, we also observed a very small number of tdTomato+ neurons in the DG hilus. These TRAPed hilar neurons were morphologically distinct from labeled granule cells (**Fig. S2a**). We found that numbers of cFos+ and tdTomato+ hilar neurons were comparable between A to A and A to B re-exposure paradigms, as well as between A to A and B to A CFC paradigms. No significant differences were observed for cFos+ tdTomato+ overlap between the paradigms (**Fig. S2a-g**), although this may be attributable to the low overall tdTomato+ cell numbers in hilus.

We (11–13) and others (9, 10) have previously shown that sleep deprivation (SD) over the hours immediately following CFC results in disrupted CFM consolidation. We confirmed these disruptive effects in the *cfos::tdTomato* mouse line. At ZT0, all mice underwent single-trial CFC in Context A, after which they were returned to their home cage for either 6 h of SD by gentle handling (followed by recovery sleep) or *ad lib* sleep (**Fig. 2a**). At ZT0 the following day, mice were returned to Context A to test CFM recall. Consistent with prior results, SD significantly reduced context-specific freezing during the CFM test (**Fig. 2b**).

**Fig. 2:**
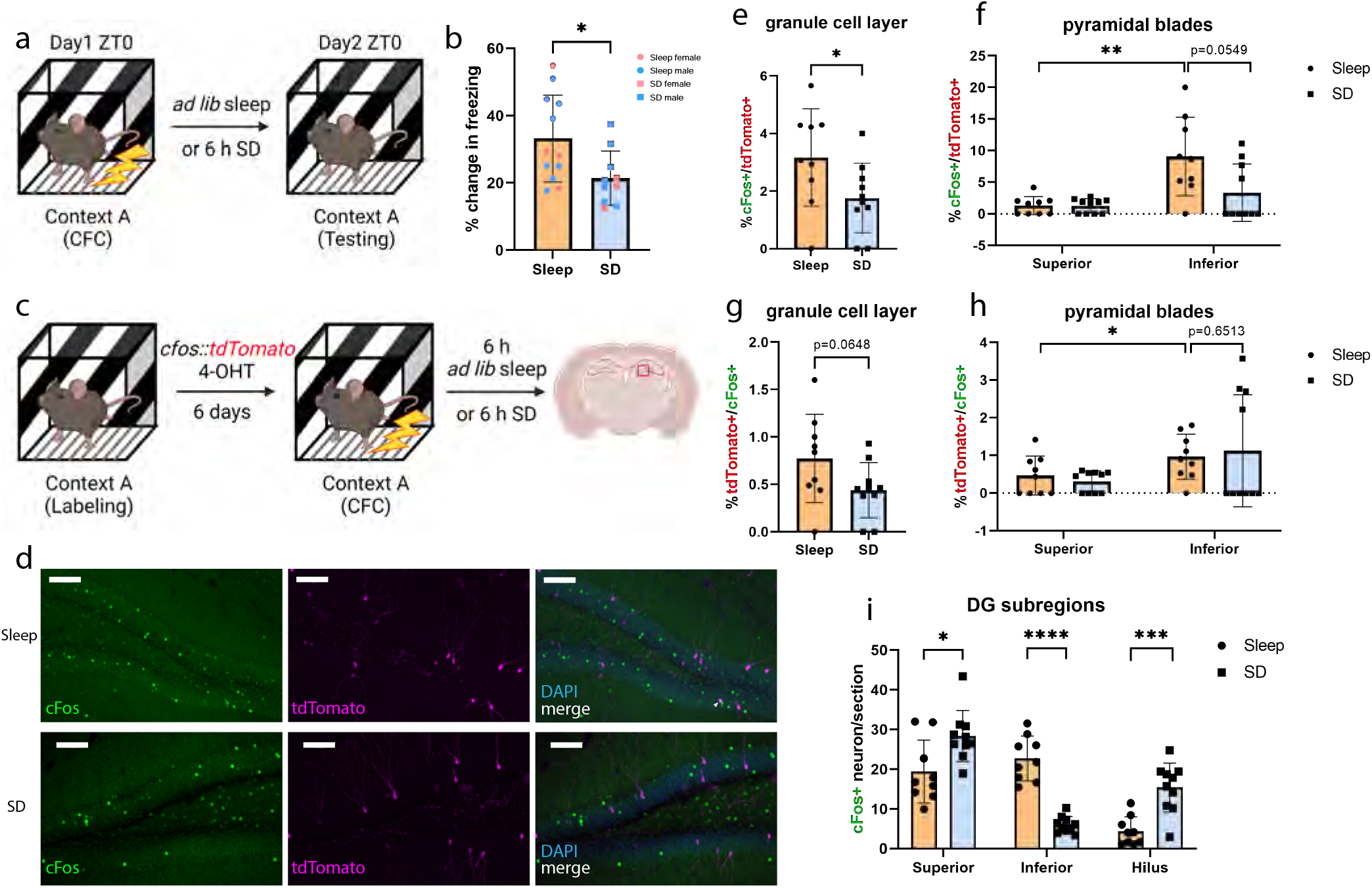
Post-learning SD disrupts reactivation of dentate gyrus engram neurons in a subregion-specific manner. **a,** Experimental procedures. Male and female mice underwent single-trial CFC at ZT0 and were either allowed *ad lib* sleep (*n* = 12, 8 males and 4 females) or underwent SD (*n* = 10, 7 males and 3 females) for the first 6 h after CFC; CFM testing occurred at ZT0 next day. **b,** CFM consolidation (measured as the change in context-dependent freezing from baseline) was significantly reduced after SD. * indicates *p* = 0.0198, Student’s t test. **c,** Male *cfos::tdtomato* mice were injected with 4-OHT following Context A exploration. 6 d later, all mice received contextual fear conditioning (CFC) in Context A and were either allowed *ad lib* sleep or were sleep deprived (SD) for 6 h prior to tissue harvest. **d,** Representative images showing overlap of tdTomato (magenta) and cFos protein (green). Examples of colocalization within a neuron is indicated with a white arrowhead. Scale bar = 100 μm. **e,** Percentage of cFos+/tdTomato+ cells in the DG granule cell layer is significantly higher following sleep (*n* = 9) than SD (*n* = 10). * indicates *p* = 0.0347, Mann Whitney test. **f,** Sleep mice had a significantly larger percentage of cFos+/tdTomato+ neurons in the inferior blade compared with the superior blade (** indicates *p* = 0.0078, Wilcoxon matched-pairs signed rank test), and a strong trend (*p* = 0.0549, Mann Whitney test) for more cFos+/tdTomato+ overlap in the inferior blade compared with the SD mice. **g,** Percentage of tdTomato+/cFos+ cells in the DG granule cell layer have a strong trend (*p* = 0.0648, Mann Whitney test) for being higher following sleep (*n* = 9) than SD (*n* = 10). **h,** Sleep mice had a significantly larger percentage of tdTomato+/cFos+ neurons in the inferior blade compared with the superior blade (* indicates *p* = 0.0391, Wilcoxon matched-pairs signed rank test). No significant difference (*p* = 0.6513, Mann Whitney test) between sleep and SD mice in the inferior blade. **i,** SD increased cFos+ cell number in the superior blade (* indicates *p* = 0.035, Mann Whitney test) and the hilus (*** indicates *p* = 0.0009, Mann Whitney test), and reduced cFos+ cell numbers in the inferior blade (**** indicates *p* < 0.0001, Mann Whitney test). All data are presented as mean ± s.d.

We next tested the effect of post-learning SD on the activity of CFC-activated engram neurons in DG in the hours following CFC (i.e., on engram neuron reactivation during consolidation). Naive *cfos::tdTomato* mice were allowed to explore Context A at ZT0, and DG context-activated neurons were labeled via 4-OHT administration. 6 days later at ZT0, mice underwent single-trial CFC in Context A, and were then returned to their home cage for either 6 h of SD or 6 h of *ad lib* sleep, after which they were perfused (**Fig. 2c**). No significant differences were observed between freely-sleeping and SD mice for total numbers of tdTomato+ or cFos+ tdTomato+ DG hilar neurons (**Fig. S2h-i**). However, consistent with observations of ensemble reactivation in the hippocampus [25] and neocortex [23], reactivation of Context A tdTomato+ DG granule cells was evident 6 h after CFC (albeit at a slightly lower rate than activation of Context A neurons during Context A CFC; **Fig. 1g-i**) (**Fig. 2d-e**). Moreover, a higher ratio of Context A tdTomato+ granule cells were cFos+ at this timepoint in freely-sleeping vs. SD mice (**Fig. 2d-e**). Surprisingly (and in contrast to effects of reexposure to context alone), this difference appeared to be driven largely by higher reactivation rates among TRAPed inferior blade granule cells during sleep (**Fig. 2f**). Freely-sleeping mice, but not SD mice, had a significantly larger proportion of cFos+ tdTomato+ granule cells in the inferior blade compared with the superior blade **(Fig. 2f,h)**. Freely-sleeping mice also showed a strong trend (*p =* 0.0549) for more cFos+ tdTomato+ granule cells in the inferior blade compared with the SD mice (**Fig. 2f**). Together, these data suggest that post-CFC sleep could promote consolidation of CFM by reactivating DG engram neurons in a subregion-specific manner that is spatially distinct from how these neurons are reactivated during waking experience, More specifically, they suggest that while superior blade DG engram neurons are preferentially activated by context during memory encoding, inferior blade engram neurons are preferentially reactivated in the hours following CFC, during sleep-dependent CFM consolidation.

Prior studies using a tTA-based system identified extensive immediate-early gene (IEG)-driven transgene expression among CA1 neurons after context exposure (26). However, consistent with prior reports using TRAP (31), we observed very sparse tdTomato expression among neurons in the CA1 and CA3 pyramidal cell layers after Context A exposure in *cfos::tdTomato* mice (**Fig. S3a-b**). In contrast to what was observed in DG, following Context A CFC, tdTomato+ cells in CA1 showed a higher level of cFos expression after post-CFC SD than following *ad lib* sleep (**Fig. S3c-d**). In comparison, there was no difference in the level of cFos expression among CA3 neurons between SD and Sleep mice (**Fig. S3e-f**). Together with data suggesting that manipulating activity among putative CA1 engram neurons does not affect CFM encoding (26), these findings suggest that reactivation of DG engram neurons in the hours following CFC might play a uniquely instructive role during sleep-dependent CFM consolidation.

### Post-learning SD selectively suppresses activity of DG inferior blade granule cells

Our data suggest subregion-specific changes in DG engram neuron reactivation in *cfos::tdTomato* mice after sleep vs. SD. Moreover, our previous findings (17, 20, 21) suggest that overall neuronal activity levels might differ between DG and other hippocampal structures as a function of sleep vs. SD. To clarify how post-CFC sleep affects network activity across hippocampal subregions, we next compared expression of IEG proteins cFos and Arc among neurons in DG, CA1, and CA3 in C57BL/6J mice following post-CFC sleep vs. SD. When single-trial CFC in Context A was followed by 6 h of SD (**Fig. 3a**), both cFos and Arc expression were significantly decreased among inferior blade granule cells (**Fig. 2i**; **Fig. 3 a-d**). However, expression in the superior blade was either unchanged or increased after SD (**Fig. 2i**, **Fig. 3c-d**), and expression of cFos in the hilus was dramatically increased after SD (**Fig. 2i**, **Fig. 3c**). Moreover, SD significantly increased cFos+ and Arc+ cell numbers in the CA3 pyramidal cell layer (**Fig. 3 e-g**), as well as relative cFos and Arc staining intensity in the CA1 pyramidal cell layer (**Fig. 3 h-j**). Taken together, these data are consistent with previous reports that SD drives alterations in DG activity that differ from those reported elsewhere in the brain (17, 20, 21) . They also suggest that following CFC, SD-mediated disruption of CFM consolidation could be caused by selectively disrupted activity (and associated engram neuron reactivation) in the DG inferior blade.

**Fig. 3:**
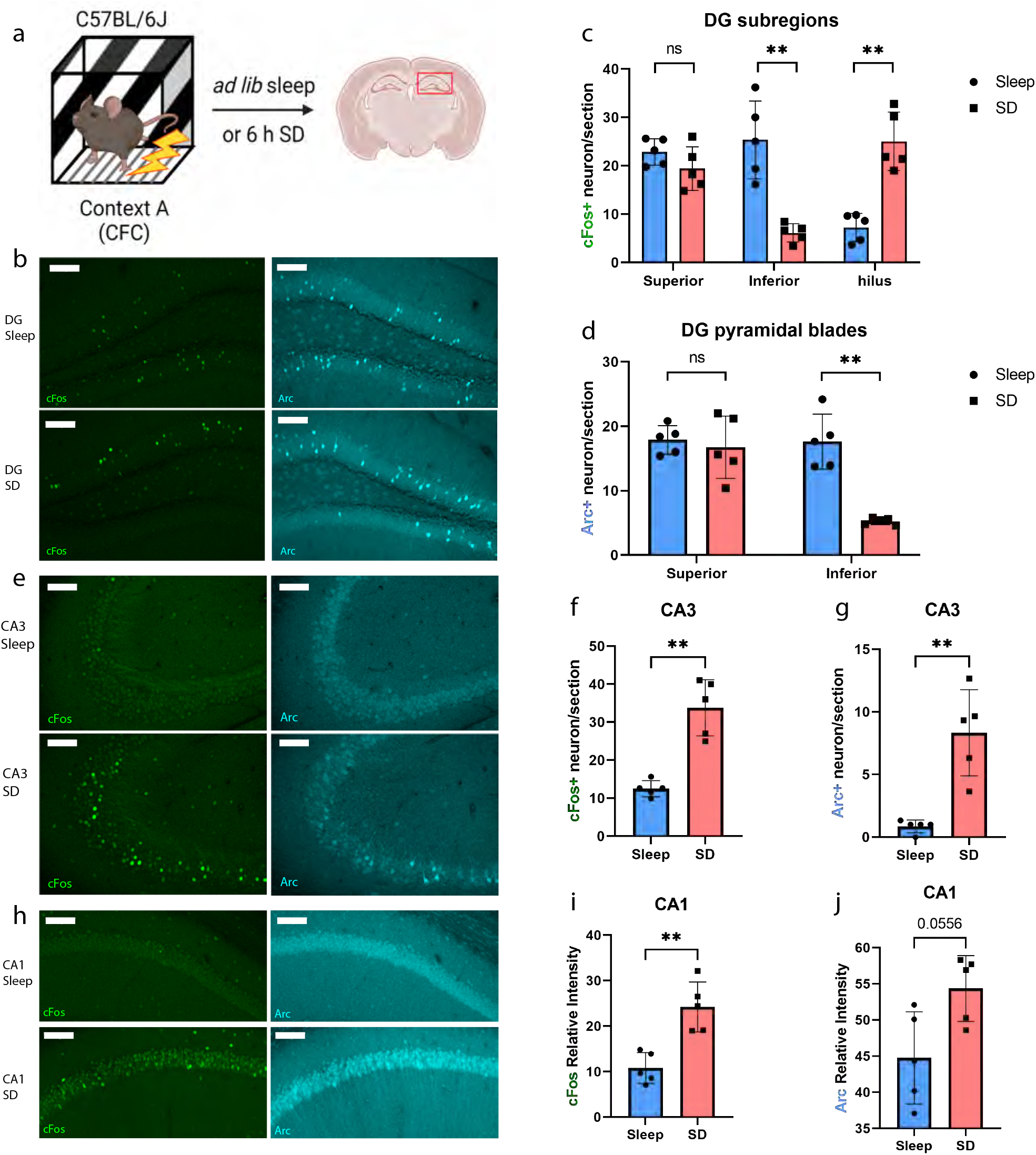
Post-learning SD selectively suppresses activity of DG inferior blade granule cells. **a,** Experimental procedure. C57BL/6J mice underwent single-trial CFC in Context A and were either allowed ad lib sleep or were sleep deprived (SD) for 6 h prior to tissue harvest. **b,** Representative images showing cFos (green) and Arc protein (cyan) following sleep or SD in the dentate gyrus. Scale bar = 100 μm. **c,** SD did not change cFos+ cell number in the superior blade in C57BL/6J mice (*p* = 0.3095, Mann Whitney test) but increased the cFos+ cell the hilus (** indicates *p* = 0.0079, Mann Whitney test), and reduced cFos+ cell number in the inferior blade (** indicates *p* = 0.0079, Mann Whitney test). **d,** SD reduced Arc+ cell numbers in the inferior blade compared with sleep. ** indicates *p* = 0.0079, Mann Whitney test. **e,** Representative images showing cFos (green) and Arc protein (cyan) following sleep or SD in the CA3. Scale bar = 100 μm. **f,** SD increased cFos+ (**indicates *p* = 0.0079, Mann Whitney test) and (**g**) Arc+ (** indicates *p* = 0.0079, Mann Whitney test) cell number in the CA3. Data are presented as mean ± s.d. **h,** Representative images showing cFos (green) and Arc protein (cyan) following sleep or SD in the CA1. Scale bar = 100 μm. **i,** SD increased cFos protein relative intensity (** indicates *p* = 0.0079, Mann Whitney test) in the CA1 pyramidal layer and (**j**) had a strong trend of increasing Arc (*p* = 0.0556, Mann Whitney test). All data are presented as mean ± s.d.

### SD causes diverse and subregion-specific alterations in hippocampal gene expression, and drives distinctive responses in the DG blades

Our preliminary findings (**Figs. 1-3**) suggest that post-CFC sleep may have differential effects on granule cell, and more specifically, engram neuron activation between the two DG blades. These subregion-specific effects may be linked to sleep-dependent CFM consolidation. We have recently shown that learning and subsequent sleep or sleep loss differentially affect ribosome-associated mRNA profiles in different hippocampal cell types (12, 21). Based on our observation of subregion-specific changes in cFos and Arc expression and engram neuron reactivation following post-CFC sleep or SD (**Figs. 2**, **3**), we speculated that learning and sleep could independently alter biosynthetic processes that impinge on synaptic plasticity in a subregion-specific manner. More specifically, we hypothesized that CFC and subsequent sleep vs. SD could differentially impact important neurobiological processes in the DG inferior blade vs. other hippocampal substructures, including the superior blade.

To test this hypothesis, we separately profiled mRNAs in various subregions within the dorsal hippocampus using NanoString GeoMx Digital Spatial Profiler (DSP). At ZT 0, mice were either left undisturbed in their home cage (HC) or underwent single-trial CFC in Context A (CFC). Over the next 6 h, mice in both the CFC and HC groups were either allowed *ad lib* sleep or underwent SD in their home cage prior to perfusion (**Fig. 4a**). This 2 × 2 experimental design structure (resulting in 4 experimental groups) allowed us to separately test for effects of learning and sleep on transcript levels within each subregion. Brain sections from each experimental group were stained with nuclear label Syto 13 to identify borders of the DG hilus, the superior and inferior blade granule cell layers of DG, and the pyramidal cell body layers of CA1 and CA3 (example regions of interest [ROIs] shown in **Fig. 4b**). From each mouse (*n* = 3-4 mice/group) either bilateral or unilateral hippocampal subregions were sampled for DSP transcript measurement (1-2 regional samples/mouse [**Table S1**], each subregion sample corresponding to one biological replicate). In total, 11508 gene targets from the mouse Whole Transcriptome Atlas (WTA) passed target filtering and were quantified for each sample. As expected, principal component analysis (PCA) revealed clear separation of CA1, CA3, DG hilus, and DG pyramidal blades’ gene expression profiles (**Fig. S4a**).

**Fig. 4:**
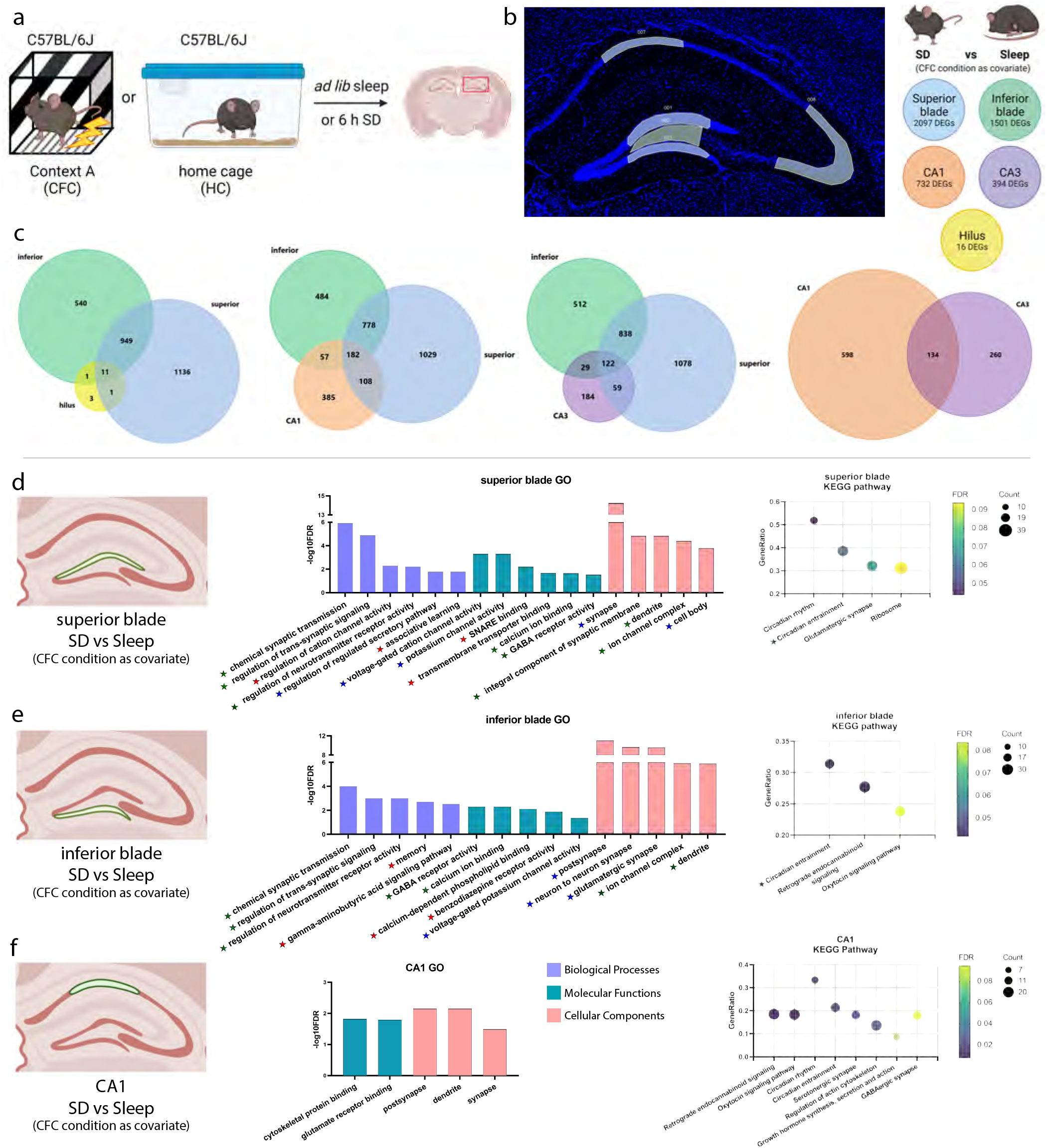
SD causes subregion-specific alterations in hippocampal gene expression. **a,** C57BL/6J mice were either left undisturbed in their home cage (HC) or underwent single-trial CFC in Context A (CFC). Over the next 6 h, mice in CFC and HC groups were either allowed ad lib sleep or underwent SD in their HC prior to perfusion. **b,** (***Left***) Representative image of (region of interest) ROI selection using NanoString’s GeoMx Digital Spatial Profiler (DSP). (***Right***) Illustration of the comparison and number of DEGs for SD vs Sleep in each hippocampal subregion. **c,** Venn diagrams reflect the number of SD vs Sleep DEGs and their overlap in the DG superior blade, inferior blade, hilus, CA1, and CA3. **d,** The most significant gene ontology terms, ranked by FDR value, and KEGG pathways enriched for transcripts altered by SD alone in the superior blade, (**e**) inferior blade, and (**f**) CA1. Red stars highlight GO terms uniquely mapped to one blade only, blue stars indicate the presence of related (parent/child) terms in both blades, and green stars highlight GO terms and KEGG pathways overrepresented in both superior and inferior blades.

We first assessed the effects of SD itself on each hippocampal subregion. For this, we included data from all mice, and used the learning condition (CFC or HC) as a covariate. This structure allowed us to 1) determine effects that are driven by SD, regardless of prior learning, and 2) make comparisons with related data sets from whole-hippocampus RNAseq data, which used a similar study design (21). Transcripts affected by SD (i.e., differentially-expressed genes [DEGs] for the SD vs. Sleep conditions; *n* = 7 and *n* = 8 mice, respectively) were then compared between hippocampal subregions (**Fig. 4b, c**). Within DG, SD significantly altered (FDR < 0.1) 2097 transcripts in the superior blade and 1501 transcripts in the inferior blade; of these, 960 altered transcripts overlapped between the two blades (**Fig. 4c**). Comparatively fewer transcripts were altered by SD in the DG hilus, although this may be due to the relatively small ROI size (in terms of contributing cell number) for the hilus when compared with granule cell layers (**Fig. S4b**).

Of the 16 SD-altered genes identified in the hilus, 11 ((*Rbm*(↓)*, Cirbp* (↓)*, Pdia6* (↑)*, Hspa5* (↑)*, Fam163b* (↓)*, Sdf2l1* (↑)*, R3hdm4* (↓), *Xbp1* (↑)*, Btf3l4* (↓)*, Dtnb* (↓), and *Sqstm1* (↓); arrows indicate increased or decreased abundance after SD) were similarly altered by SD in both the superior and inferior blades (**Fig. 4c**). For CA1 and CA3 pyramidal layers, 732 genes and 394 SD DEGs, respectively, were identified; of these, only 134 were similarly altered by SD in both subregions (**Fig. 4c**). While many SD DEGs overlapped between different hippocampal subregions (with the largest overlap between superior and inferior blades), the majority of SD DEGs was unique to individual subregions (**Fig. S5a-b**). This was true for transcripts that were either upregulated or downregulated after SD (**Fig. S5c-d**). 53 DEGs were altered by SD across CA1, CA3, DG superior and inferior blades, and these were consistently either upregulated or downregulated by SD across all four of these subregions (**Fig. S5e**). Somewhat surprisingly, only five transcripts in total (*Rbm3* (↓)*, Pdia6* (↑)*, Hspa5* (↑)*, Sdf2l1* (↑), and *Dtnb* (↓)) were consistently altered by SD across all five hippocampal subregions measured. These pan-hippocampal transcript changes included transcripts encoding components of the ER chaperone complex **(Fig. S6)**, consistent with previous findings (33) that SD activates the ER stress response, both in the hippocampus (12, 16) and elsewhere in the brain (34).

To better understand cellular mechanisms affected by these subregion-specific transcriptomic changes, we used gene ontology (GO) classifiers of biological process, molecular functions, and cellular component annotation of SD-altered transcripts in each subregion (**Fig. 4 d-f**). We first compared the two DG blades’ responses to SD against each other. Several GO terms were either overrepresented among transcripts altered by SD in both the DG superior and inferior blades (marked with green stars in **Fig. 4 d-e**), or shared parent/child terms for transcripts altered by SD in both blades (blue stars). These included transcripts encoding synaptic and dendritic components of neurons, and those involved in regulation of GABA receptors, voltage-gated channels, and synaptic transmission. A few GO terms were uniquely altered by SD in each blade (red stars). For example, transcripts annotated as mediators of memory (GO:0007613; smallest common denominator-corrected *p_adj_* = 0.002) were overrepresented among transcripts altered by SD in the inferior blade only. In contrast, transcripts annotated as mediators of associative learning (GO:0008306; *p_adj_* = 0.016) were selectively altered by SD in the superior blade. While these two biological processes are often linked together conceptually, the number of transcripts present in both annotation categories was only 318, from a total of 3,140 and 1,545, respectively. Critically, however, in both cases, the vast majority of SD DEGs mapped to these processes were downregulated after SD. We also compared KEGG database pathway mapping for SD-altered transcripts from superior and inferior blade, to identify cellular pathways differentially affected by SD in the two blades. The circadian entrainment (KEGG: 04713) pathway was affected by SD in both DG superior (FDR-corrected *p_adj_* = 0.060) and inferior (*p_adj_* = 0.042) blades. In the superior blade only, circadian rhythm (KEGG: 04710; *p_adj_* = 0.044), glutamatergic synapse (KEGG: 04724; *p_adj_* = 0.070), and ribosome (KEGG: 03010; *p_adj_* = 0.094) pathways were overrepresented among SD-altered transcripts. DEGs in the glutamatergic synapse pathway were primarily downregulated, with a few notable (and likely physiologically-important) exceptions including *Grm1*, *Grm2*, *Cacna1c*, *Gria1*, and *Grin2a*, which were all upregulated after SD (**Fig. S7a**). Interestingly, among ribosome pathway DEGs from the superior blade, SD upregulated those annotated as mitochondrial ribosome components, while downregulating those annotated as non-mitochondrial ribosome components, with no exceptions (**Fig. S7a**). This suggests strong and opposing effects of SD on mitochondrial vs. non-mitochondrial biogenesis, and supports recent work suggesting effects of SD on mitochondrial structure and function in neurons (35). In contrast to pathways identified in superior blade, inferior blade SD-altered transcripts overrepresented retrograde endocannabinoid signaling (KEGG: 04723; *p_adj_* = 0.042) and oxytocin signaling (KEGG: 04921; *p_adj_* = 0.084) pathway components. For both pathways, DEGs were generally downregulated after SD, with only a few exceptions (**Fig. S7b**).

For comparison with the DG blades, we identified GO and KEGG pathways overrepresented among CA1 SD-altered transcripts (**Fig. 4f**). Several pathways identified in this analysis were also mapped to SD DEGs from the DG blades – i.e., retrograde endocannabinoid signaling (KEGG: 04723; *p_adj_* = 0.008), oxytocin signaling (KEGG: 04921; *p_adj_* = 0.008), circadian rhythm (KEGG: 04710; *p_adj_* = 0.019), and circadian entrainment (KEGG: 04713; *p_adj_* = 0.019). As was true in DG, transcripts in these pathways were generally downregulated after SD, with the exception of circadian rhythm, where most transcripts were upregulated. Additional KEGG-annotated pathways with transcripts enriched among CA1 SD DEGs included serotonergic synapse (where serotonin receptor-encoding transcripts were generally upregulated after SD; KEGG: 04726; *p_adj_* = 0.027), regulation of actin cytoskeleton (where transcripts were generally downregulated after SD; KEGG: 04810; *p_adj_* = 0.036), growth hormone synthesis, secretion and action (KEGG: 04935; *p_adj_* = 0.084), and GABAergic synapse (where most transcripts were downregulated after SD; KEGG: 04727; *p_adj_* = 0.095) (**Fig. S7c**).

Circadian entrainment (KEGG: 04713) was the only KEGG pathway mapped for SD DEGs in all three subregions (DG superior blade, inferior blade, and CA1). Pathway diagrams of circadian entrainment showed a consistent SD-induced negative perturbation of calcium-calmodulin kinase (CaMKII) and mitogen-activated protein kinase (MAPK) signaling pathway components across three subregions, which is consistent with prior studies showing that SD attenuates kinase signaling and CREB-mediated gene transcription (10, 21, 36). Surprisingly, although MAPK and CREB signaling components upstream of clock genes were negatively regulated by SD in all three regions, clock gene and IEG transcripts were differentially perturbed across the three regions (**Fig. S8**). These findings, consistent with our observations in **Figs. 2** and **3**, indicate that multiple regulatory mechanisms drive differential SD-driven IEG expression changes across the hippocampal subregions.

To further characterize cellular pathways regulating SD-mediated changes, we performed predicted upstream regulator analysis of SD DEGs within each subregion (**Table S2**). For superior blade DEGs, this analysis predicted inhibition of upstream regulators encoded by *Cask*, *Lin7b*, *Lin7c*, *Lin7a*, *Apba1*, and *Unc13b* transcripts (FDR = 0.007), due to downregulation of *Ppfia4*, *Cplx1*, *Tspoap1*, *Rab3a*, *Stx1a*, *Syn1*, *Snap25*, and *Ppfia2* transcripts. Collectively, these downregulated transcripts encode multiple receptor tyrosine phosphatases and active zone vesicular trafficking and release regulating proteins after SD. The expression for two of these upstream regulators, *Lin7b* (encoding lin-7 homolog B, crumbs cell polarity complex component – which normally coordinates their activity in the presynaptic terminal) and *Apba1* (encoding amyloid beta precursor protein binding family A, member 1), were measured to be downregulated by SD (**Table S2**). These findings suggest that presynaptic biogenesis may be decreased within the DG superior blade after a period of SD.

Similar analysis of SD DEGs within CA1 suggested that regulators encoded by *Rab18* (encodes a member of the Ras-related small GTPases, which regulate membrane trafficking in organelles and transport vesicles), *Rab3gap2,* and *Rab3gap1* (RAB3 GTPase activating protein subunit 1 and 2) were predicted as inhibited upstream regulators, and miRNAs *mmu-miR-27a-3p* and *mmu-miR-27b-3p* were predicted as activated miRNAs.

### DG transcriptional responses to SD can oppose those in CA1 and CA3, and suggest selective disruption of DG inferior blade activity

For several SD-altered transcripts, expression levels changed in opposite directions when comparing the two DG blades with other hippocampal subregions. For example, the expression of *Arc* was upregulated by SD in hilus but downregulated by SD in the inferior blade **(Fig. 5b)**. *Zmat3* (encoding a zinc finger domain transcription factor)*, C1ql3* (encoding an extracellular regulator of excitatory synapses), and *Sf3b6* (encoding an mRNA splicing factor) were upregulated by SD in superior blade but downregulated in CA1; *Per1, Kctd12* (encoding an auxiliary subunit of GABA-B receptors)*, D830031N03Rik* (a.k.a., *Macf1;* microtubule-actin crosslinking factor 1), ephrin receptor-encoding *Epha10,* small GTPase-encoding *Rasl11b, Ets2* (encoding a telomerase-regulating transcription factor)*, Mbnl1* (encoding a pre-mRNA splicing factor)*, Gga3* (encoding a trans-Golgi network sorting and trafficking protein), and *Ier5* (which encodes an immediate early response protein that may mediate transcriptional responses to heat shock) were upregulated by SD in CA1 but downregulated in superior blade **(Fig. 5c)**. This suggests that nuclear and cytoplasmic processes, and regulators of both glutamatergic and GABAergic synapses, are differentially regulated between CA1 and superior blade in response to SD. Chromatin remodeling factor *Rbbp7, Lrrc7* (encoding a protein required for glutamate receptor localization and synaptic plasticity), postsynaptic scaffold encoding *Dlgap1, Parp1* (encoding a chromatin-associated enzyme critical for DNA damage repair)*, Rchy1* (encoding an E3 ubiquitin ligase), and *Tacc2* (encoding a regulator of nuclear structure) were upregulated by SD in superior blade but downregulated in CA3. These differences may reflect differential disruption of nuclear and cytoplasmic functions between the two regions, resulting in increased engagement of nuclear and cytoplasmic quality control mechanisms in the superior blade. On the other hand, *Chrd* (encoding secreted factor chordin), *Car2* (encoding carbonic anhydrase), *Apc2* (encoding a negative regulator of the Wnt signaling pathway), *Rell2* (encoding a collagen-binding activator of the MAPK pathway), and *Iyd* (encoding a tyrosine deiodinating enzyme) were upregulated by SD in CA3 but downregulated in superior blade **(Fig. 5e)**.

**Fig. 5:**
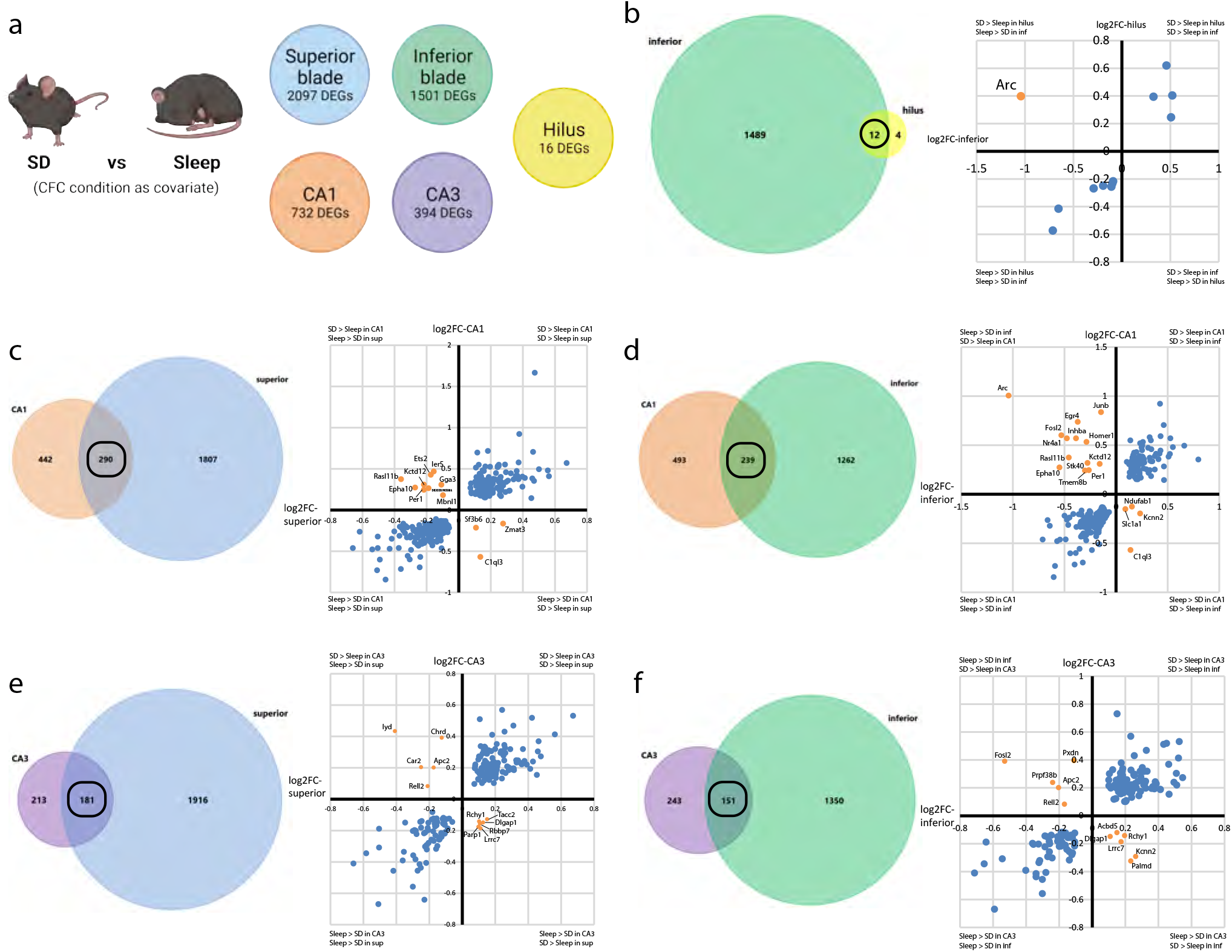
Select SD-altered transcripts’ levels were altered in opposite directions across hippocampal subregions. **a,** Illustration of the comparison and number of DEGs for SD vs Sleep in each hippocampal subregion. **b,** Expression level of the 12 overlapping DEGs between inferior blade and hilus. Blue dots represent SD-altered DEGs that are consistently enriched in one of the two subregions. Orange dot represents the gene (*Arc*) that is upregulated by SD in hilus but downregulated by SD in DG inferior blade. **c,** Expression level of the 290 overlapping DEGs between superior blade and CA1. 3 DEGs were upregulated by SD in DG superior blade but downregulated in CA1 and 9 were upregulated by SD in CA1 but downregulated in superior blade. **d,** Expression level of the 239 overlapping DEGs between inferior blade and CA1. Of these, 4 were upregulated by SD in the inferior blade but downregulated in CA1 and 13 were upregulated by SD in CA1 but downregulated in the inferior blade. **e,** Expression level of the 181 overlapping DEGs between superior blade and CA3. 6 were upregulated by SD in the superior blade but downregulated in CA3 and 5 were upregulated by SD in CA3 but downregulated in the superior blade. **f,** Expression level of the 151 overlapping DEGs between inferior blade and CA3. 6 were upregulated by SD in the inferior blade but downregulated in CA3 and 5 were upregulated by SD in CA3 but downregulated in the inferior blade. Blue dots represent SD-altered DEGs that are consistently enriched in one of the two subregions. Orange dots represent DEGs that are regulated in opposite directions by SD in the subregions.

Consistent with our immunohistochemical data **(Fig. 3)**, expression of many IEGs, including *Arc, Fosl2, Homer1, Nr4a1, Junb, and Egr4*, was differentially affected by SD in CA1 vs. inferior blade, indicating opposite changes in neuronal activation patterns during SD in these structures. Similar to the superior blade, expression of *Epha10, Per1, Rasl11b,* and *Kctd12* were decreased after SD in the inferior blade, and simultaneously increased by SD in CA1. In addition, *Tmem8b* (encoding a cell matrix adhesion protein)*, Stk40* (encoding a serine-threonine protein kinase)*, Inhba* (encoding a protein involved in hormone secretion) were downregulated by SD in the inferior blade, and upregulated by SD in CA1. On the other hand, *Ndufab1* (encoding part of mitochondrial respiratory complex 1), *Kcnn2* (encoding a calcium-activated potassium channel that prolongs action potential afterhyperpolarization, and would be predicted to reduce neuronal firing rates), *C1ql3* (encoding an extracellular regulator of excitatory synapses), and *Slc1a1* (encoding a high-affinity glutamate transporter) were upregulated by SD in the inferior blade but downregulated in CA1 **(Fig. 5d)**. Taken together, these transcriptomic changes are consistent with an overall reduction of neuronal activity during SD that occurs selectively in the DG inferior blade. *Kcnn2, Lrrc7, Dlgap1, Rchy1, Palmd* (encoding paralemmin-like protein, which is predicted to be involved in dendritic remodeling), and *Acbd5* (encoding an Acyl-CoA binding protein) were upregulated by SD in inferior blade but downregulated in CA3, in large part mirroring differences between superior blade and CA3. *Prpf38b* (encoding a pre-mRNA splicing factor), *Pxdn (*encoding a peroxidase involved in extracellular matrix formation), AP-1 transcription factor subunit *Fosl2, Apc2* (a microtubule stabilizing factor, and putative negative regulator of Wnt signaling), and *Rell2* were upregulated by SD in CA3 but downregulated in the inferior blade (**Fig. 5f**). These differences suggest that transcripts associated with neuronal activity and synaptic and structural plasticity are simultaneously upregulated in CA3 and downregulated in the inferior blade after SD.

In contrast to the differential regulation of individual transcripts between the DG blades and CA1/3 subregions after SD, no transcripts were regulated in different directions when comparing the hilus, CA1, and CA3 DEGs against each other (**Fig. S9**). Together, these data support the conclusions that: 1) SD differentially affects granule cells in the two DG blades vs. pyramidal cells of CA3 and CA1, and 2) selective effects of SD on the inferior blade’s transcript profile may reflect the selective suppression of activity in the inferior blade during SD.

### Transcriptomic differences between superior and inferior DG blades are altered by SD vs. sleep

Because we observed differences in engram neuron context selectivity and sleep-associated reactivation between the DG inferior blade and superior blades (**Figs. 1** and **2**), we directly compared how gene expression differed between the two blades (Inferior vs. Superior DEGs) after sleep vs. SD, again using prior learning condition (CFC/HC) as a covariate. 580 genes were differentially expressed between inferior and superior blade in freely-sleeping mice (Sleep Inferior vs. Sleep Superior), while 750 genes were differentially expressed between the blades in SD mice (SD Inferior vs. SD Superior). 251 Inferior vs. Superior DEGs overlapped between Sleep and SD conditions (**Fig. 6a**), of which 249 were consistently expressed at a higher level in either the inferior or superior blades (**Fig. 6b**). For example, *Penk, Fst, Pmepa1, Npy, Col6a1,* and 156 other DEGs were consistently enriched in the superior blade while *Sema5b, Lhx9, Myo5b, Gpc3* and 84 additional DEGs were consistently enriched in the inferior blade (**Fig. 6b**); these likely represent differences in cellular constituents between the blades, some of which have been previously characterized. For example, several DEGs we identified as enriched in the superior blade, including *Penk, Rgs4*, *Col6a1*, and *Nefm*, may be attributed to a recently-identified subcluster of *Penk*-expressing granule cells located in the DG superior blade (32, 37). These differentially-expressed transcripts thus likely reflect true (constitutive) genetic differences between the two blades.

**Fig. 6:**
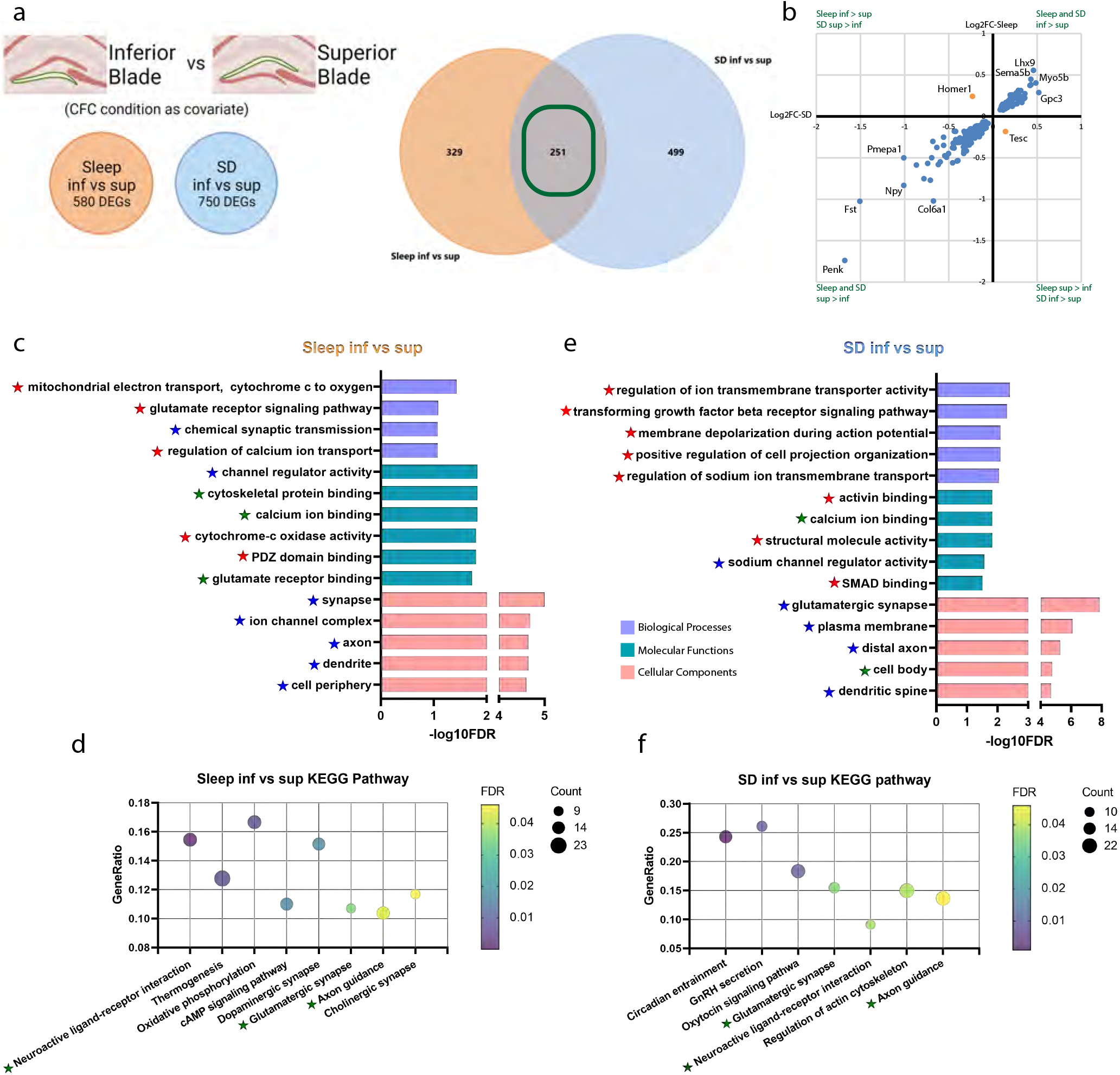
Transcriptomic differences between superior and inferior DG blades. **a,** (***Left***) Illustration of the inferior blade vs superior blade comparison and the number of DEGs under either Sleep or SD condition. (***Right***) Venn diagram reflects the overlap (251 transcripts) of inferior vs superior blade DEGs under Sleep or SD condition. **b,** 249 (blue dots) of the 251 DEGs were consistently expressed at a higher level in either the inferior or superior blades. *Homer1* was enriched in the superior blade in SD mice but enriched in the inferior blade in Sleep mice, while *Tesc* was enriched in the inferior blade in SD mice but enriched in the superior blade in Sleep mice. **c, e,** The most significant gene ontology terms and **d, f** KEGG pathways - ranked by FDR values - mapped for inferior vs superior DEGs in (**c, d)** Sleep mice (**e, f**) and SD mice. Red stars highlight GO terms uniquely mapped under either Sleep or SD condition, blue stars indicate the presence of parent/child terms for both conditions, and green stars highlight GO terms and KEGG pathways overrepresented in both Sleep and SD conditions.

A few Inferior vs. Superior DEGs showed very dramatic alterations as a function of Sleep vs. SD. For example, consistent with our immunohistochemical results (**Fig. 3**), *Fos* and *Arc* showed higher expression levels in the superior blade than the inferior blade after SD, but did not differ between the blades in freely-sleeping mice. Interestingly, some transcripts were differentially enriched in one blade vs. the other as a function of sleep condition. For example, *Homer1* (encoding an activity-regulated protein involved in synaptic growth and hippocampal plasticity) (38, 39) expression was higher in the superior blade after SD, but with *ad lib* sleep, its expression was higher in the inferior blade (**Fig. 6b**). On the other hand, *Tesc* (encoding a calcium-binding protein implicated in dendritic growth and neuronal survival) (40) expression was higher in the inferior blade with SD, but with *ad lib* sleep, its expression was higher in the superior blade (**Fig. 6b**).

We performed GO and biological pathway analysis on Inferior vs. Superior DEGs to further explore how the two blades’ function differed under Sleep and SD conditions (**Fig. 6 c-f**). Most cellular component terms had partially overlapping parent/child terms mapped under either the Sleep or the SD condition (blue stars), while biological process and molecular function had more terms uniquely mapped under either the Sleep or the SD condition, with no direct parent/child relationships (red stars). For example, Sleep was associated with Inferior vs. Superior DEGs enriched for PDZ domain binding molecular function (GO:0030165; smallest common denominator-corrected *p_adj_* = 0.016) – typically membrane-associated synaptic signaling molecules. Within this molecular class, after *ad lib* sleep, the inferior blade had lower expression of some transcripts, including *Cit* (encoding citron, an actin cytoskeleton-associated serine/threonine kinase activating protein), *Cntnap2*, and MAGUK family members *Mpp3* and *Dlg3*, but higher expression of others, including glutamate receptor components *Grid2*, *Grik2*, and *Gria2*, AMPA receptor regulator *Shisa9*, and voltage-gated calcium channel subunit *Cacng3*. Several pathways were identified using KEGG analysis as differentially regulated between the two DG blades after sleep, including glutamatergic synapse (KEGG: 04724; FDR-corrected *p_adj_* = 0.035), neuroactive ligand-receptor interaction (KEGG: 04080; *p_adj_* = 8.171e-5), axon guidance (KEGG: 04360; *p_adj_* = 0.043), thermogenesis (KEGG: 04714; *p_adj_* = 0.005), oxidative phosphorylation (KEGG: 00190; *p_adj_* = 0.005), cAMP signaling pathway (KEGG: 04024; *p_adj_* = 0.017), dopaminergic synapse (KEGG: 04728; *p_adj_* = 0.017), and cholinergic synapse (KEGG: 04725; *p_adj_* = 0.046). For all of these pathways, most or all DEG components were expressed at significantly lower levels in the inferior blade, relative to the superior blade, after sleep.

In SD mice, Inferior vs. Superior DEGs enriched for components of the circadian entrainment (KEGG: 04713; *p_adj_* = 9.796e-4), GnRH secretion (KEGG: 04929; *p_adj_* = 0.011), oxytocin signaling (KEGG: 04921; *p_adj_* = 0.011), glutamatergic synapse (KEGG: 04724; *p_adj_* = 0.036), neuroactive ligand-receptor interaction (KEGG: 04080; *p_adj_* = 0.038), regulation of actin cytoskeleton (KEGG: 04810; *p_adj_* = 0.039), and axon guidance (KEGG: 04360; *p_adj_* = 0.046) pathways. Only three of these pathways also mapped to Inferior vs. Superior DEGs in the freely-sleeping mice (green stars; **Fig. 6d, f**). Interestingly, Inferior vs. Superior DEGs enriched for regulation of actin cytoskeleton only in the SD condition. The vast majority of DEGs in this pathway were expressed at lower levels in the inferior blade vs. superior blade after SD, including *Mylk3* (encoding myosin light chain kinase 3), *Myh10* (myosin heavy polypeptide 10, involved in postsynaptic cytoskeleton reorganization), *Arpc2*, *3*, and *4* (actin related protein 2/3 complex, subunits 2, 3, and 4; subunit 2 positively regulates actin polymerization and localize to glutamatergic synapses), *Actr2* (Arp2 actin-related peptide 2, which localizes to the actin cap), *Arhgef4* (Rho guanine nucleotide exchange factor 4, thought to be involved in filopodium assembly), *Pfn1* and *Pfn2* (profilin-1 and profilin-2, regulators that bind to actin and affect the structure of the cytoskeleton), and *Pak1* (P21/RAC1 Activated Kinase 1, a cytoskeletal regulator of cell motility and morphology) (**Fig. S10**). The selective downregulation of these pathway components in the inferior blade after SD likely results in disruption of structural plasticity in inferior blade neurons. This idea is supported by recent data showing that SD selectively reduces dendritic spine density in inferior blade granule cells, while leaving superior blade granule cells relatively unaffected (41).

Upstream regulator analysis (**Table S2**) further predicted that after SD (but not after *ad lib* sleep), upstream regulators *Exosc10, Mtrex, C1d,* and *Bud23* would be inhibited in the inferior blade. This is based on downstream downregulated expression of many transcripts encoding ribosomal protein components (*Rps2,6,7,9,* and *14,* and *Rpl3, 4, 5, 19, 27a,* and *38*) in the inferior blade relative to the superior blade after SD. These transcripts are known to be directly transcriptionally coregulated by two nucleolar factors, *Exosc10* (exosome component 10) and *Mtrex* (Mtr4 exosome RNA helicase), although neither *Exosc10* nor *Mtrex* transcript was differentially expressed between the two DG blades after SD. Another upstream regulator predicted to be suppressed in the inferior blade after SD was *Wasl* (WASP like actin nucleation promoting factor). Consistent with this prediction, a number of transcripts encoding actin-regulating factors (*Arpc2, 3,* and *4*, and *Atcr2*) were all selectively downregulated in the inferior blade relative to the superior blade after SD (but not after *ad lib* sleep), although expression levels for *Wasl* itself did not differ between the blades after SD.

Because *de novo* protein synthesis and actin cytoskeleton rearrangement are thought to be critical for synaptic plasticity and memory consolidation, the selective disruption of these two cellular pathways in the inferior blade after SD may lead to SD-mediated memory consolidation deficits. Taken together, these analyses identify numerous functional differences between the superior and inferior blades which appear specifically under SD conditions. These SD-driven changes are likely to disrupt synaptic and structural plasticity in inferior blade granule cells, and are consistent with the selective disruption of engram neuron reactivation in the inferior blade during SD. These transcriptomic differences may either represent either an upstream cause, or a downstream consequence, of this subregion-specific activity disruption.

### CFC-induced transcriptomic effects in the hours following learning are restricted to the hippocampal DG

Recent findings using translating ribosome affinity purification sequencing have revealed that transcriptomic effects of CFC vary with cell type in the hippocampus, and are almost exclusively restricted to ribosomes associated with neuronal cell and organellar membranes (12). To understand how CFC affects transcripts in different hippocampal subregions, we quantified transcriptomic effects of learning (comparing CFC vs. HC) using data from all mice, using sleep condition (Sleep or SD) as a covariate (**Fig. 7a**). In dramatic contrast to the more widespread effects of Sleep vs. SD (**Fig. 4c**), CFC itself had no significant effects on transcripts measured 6 h later in CA3 or CA1 pyramidal cell layers, or in the DG hilus. CFC altered the most transcripts (784 CFC vs. HC DEGs) within the DG superior blade. We identified two cellular component GO terms -synaptic membrane (GO:0097060, smallest common denominator-corrected *p_adj_* = 0.016) and GABA receptor complex (GO:1902710, *p_adj_* = 0.045) - that had constituent transcripts significantly enriched among DEGs altered in the superior blade by CFC (**Fig. 7c**). 45 DEGs mapped to the synaptic membrane term. Transcripts downregulated in the superior blade after CFC included GABAA receptor subunit-encoding genes *Gabra3, Gabrg3,* and *Gabra2, Rims3* (encoding a presynaptic regulator of exocytosis), *Dcc* (encoding a protein with pre and postsynaptic roles in excitatory synapse plasticity), *Lrfn5* (encoding a cell surface regulator of synapse assembly), *Ache* (encoding acetylcholinesterase), *L1cam* (encoding an presynaptic cell surface adhesion regulator), *Trappc4* (trafficking protein particle complex 4, which is involved in dendrite development and endosome regulation), *Kcnc2* (encoding voltage-gated potassium channel Kv3.2), *Tenm2* (teneurin transmembrane protein 2), *Cntnap4* (contactin associated protein-like 4, a regulator of GABAA receptor localization and transmission), and *Ank1* (which links synaptic membrane proteins to the spectrin cytoskeleton) (**Fig. S11c**). Many more transcripts in the GO term were upregulated in the superior blade after CFC, including those encoding both metabotropic (*Grm2*) and ionotropic (*Gria1*, *Gria2*, *Grin2a*) glutamate receptor subunits, two cornichon family members (*Cnih2* and *Cnih3,* auxiliary AMPA receptor subunits that promote receptor transmission), components of the presynaptic vesicle release machinery (*Erc2*, *Stx3, Syt7*), potassium (*Kcna1*, encoding voltage gated delayed rectifier Kv1.1, and *Kcnj6*, encoding presynaptic inward rectifier GIRK2) and calcium channels (*Cacna1h*, encoding T-type voltage gated Ca(v) 3.2), metabotropic GABA receptor 2 (*Gabbr2*), regulator of postsynaptic receptor internalization synaptojanin (*Synj1*), postsynaptic MAGUK scaffold protein discs large 2 (*Dlg2*), inositol 1,4,5-trisphosphate receptor 1 (*Itpr1*), *Dbn1* (encoding drebrin1, a postsynaptic actin and profilin interacting protein that positively regulates dendritic spine morphogenesis and glutamatergic receptor localization), *Rph3a* (rabphilin 3a, which retains NMDA receptors at the postsynaptic density), and *Ank3* (which anchors transmembrane proteins at both synaptic structures and the axon initial segment). CFC also upregulated transcripts encoding synaptic membrane proteins with less well-characterized function, but which have been identified as risk genes for autism in genome-wide screens (*Nipsnap1*, *Iqsec2, Adgrl1*, *Epb41l1,* and *Pten*) (**Fig. S11d**). Six CFC vs. HC DEGs were also associated with the GABA receptor complex term, including *Gabra3*, *Gabrg3*, *Gabra5*, *Gabbr2*, *Gabra2*, *Gabrb3* (**Fig. 7d**). The overall profile of transcript changes - with downregulated transcripts related to GABAergic signaling and upregulated transcripts related to structural remodeling and glutamatergic signaling - suggests that excitatory-inhibitory balance may change in the superior blade following learning. Together, these data support the same conclusion as our prior work (12) – that CFC strongly affects membrane-associated and synaptic components of neurons – and indicates that these post-learning synaptic and structural changes are present for many hours post-CFC across the DG superior blade.

**Fig. 7:**
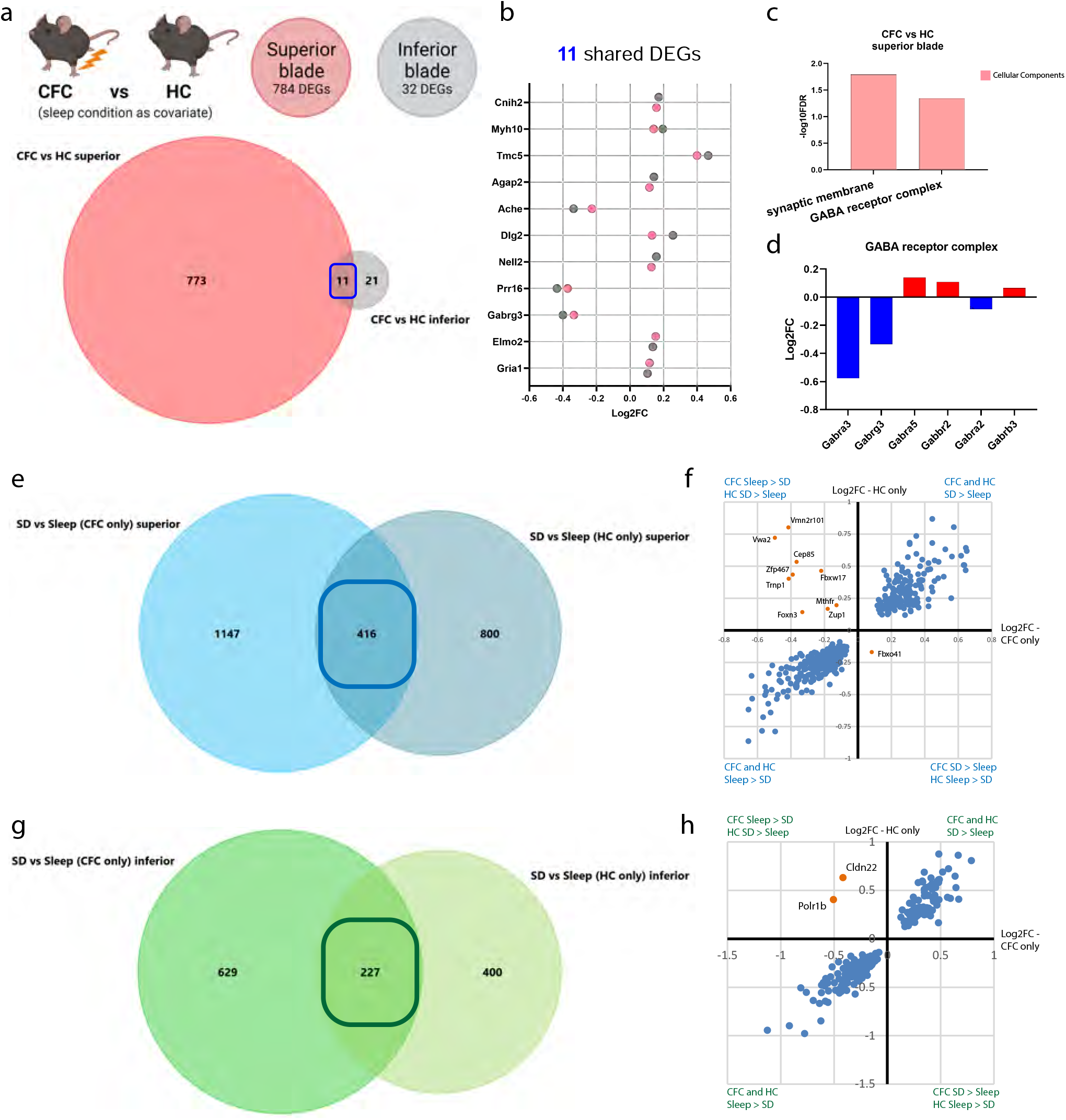
Learning (CFC)-induced transcriptomic effects are restricted to the hippocampal DG. **a,** Illustration of the CFC vs HC comparison and the number of DEGs in superior blade and inferior blade. There are no DEGs found in the CA1, CA3, and hilus for the CFC vs HC comparison. Venn diagram reflects the overlap (11 transcripts) for CFC vs HC. **b,** CFC altered 11 transcripts significantly in both superior (red) and inferior blade (gray). **c,** Significant GO terms mapped for the DEGs altered by CFC in the superior blade. **d,** DEGs altered by CFC mapped for the GABA receptor complex GO term. **e,** In mice that received CFC, 1563 transcripts were altered by subsequent SD in the superior blade while in HC mice, the number was 1216. 416 of them overlapped. **f,** Expression level of the 416 overlapping SD-altered DEGs between CFC-only and HC-only mice. Blue dots represent transcripts that are consistently enriched in the two groups following either sleep or SD. Orange dots represent genes that are regulated in opposite directions by SD in the two groups of mice. **g,** In CFC mice, 856 transcripts were altered by subsequent SD in the inferior blade while in HC mice, the number was 627. 227 of them overlapped. **h,** Expression level of the 227 overlapping SD-altered DEGs between CFC-only and HC-only mice. *Polr1b* and *Cldn22* (orange dots) were upregulated by SD in HC mice, but downregulated by SD in CFC mice.

In comparison with the relatively large number of CFC vs. HC DEGs in the DG superior blade, only 32 transcripts were altered by CFC in the inferior blade (**Fig. 7a**). Of these, 11 transcripts (*Cnih2* (↑)*, Myh10* (↑)*, Tmc5* (↑)*, Agap2* (↑)*, Ache* (↓)*, Dlg2* (↑)*, Nell2* (↑)*, Prr16* (↓)*, Gabrg3* (↓)*, Elmo2* (↑), and *Gria1* (↑)) were similarly altered in both superior and inferior blades after CFC vs. HC (**Fig. 7b**). While fewer in number, this set of transcript changes is nonetheless consistent with effects on synaptic plasticity and changing excitatory-inhibitory balance, which may be present in both DG blades following learning.

To further characterize the possible interactions between learning and subsequent sleep or SD, we next assessed the effects of SD separately in the DG blades of CFC and HC mice. In CFC mice (*n* = 4 for CFC-Sleep and *n* = 4 for CFC-SD), SD significantly altered expression of 1563 and 856 transcripts in the superior and inferior blades, respectively, while in HC mice (*n* = 4 for HC-Sleep and *n* = 3 for HC-SD), SD altered 1216 and 627 transcripts in the superior and inferior blades, respectively **(Fig. 7e, g)**. Of these, only 416 SD DEGs overlapped between CFC and HC mice in the superior blade, and 10 of those 416 overlapping transcripts were differentially regulated based on prior learning (**Fig. 7f**). These included *Vmn2r101* (encoding a putative g protein-coupled receptor), *Vwa2* (encoding an extracellular matrix protein), and a number of factors involved in DNA regulation: *Cep85* (centrosomal protein 85)*, Zfp467* (zinc finger protein 467)*, Trnp1* (TMF1 regulated nuclear protein 1, a putative euchromatin DNA-binding factor), *Fbxw17* (F-box and WD-40 domain protein 17), *Foxn3* (a member of the forkhead/winged helix transcription factor family)*, Mthfr* (methylenetetrahydrofolate reductase), and *Zup1* (zinc finger containing ubiquitin peptidase 1, thought to be involved in regulating DNA repair), which were upregulated by SD in HC mice but downregulated by SD in CFC mice, and *Fbxo41* (encoding a neuron-specific F-box protein), which was upregulated by SD in CFC mice but downregulated in HC mice (**Fig. 7f**). Similarly, in the inferior blade, only 227 SD-altered transcripts overlapped between CFC and HC mice. *Polr1b* (RNA polymerase I polypeptide B) and *Cldn22* (claudin 22) were upregulated by SD in HC mice, but downregulated by SD in CFC mice (**Fig. 7h**). While the function of these differential transcriptomic responses to SD is a matter of speculation, one possibility is that DNA repair and transcription regulatory mechanisms are differentially engaged by SD, depending on whether a memory has been recently encoded. What is clear is that for the above genes, the effects of sleep loss are dependent on prior learning experiences.

### SD and learning differentially affect protein and phosphoprotein abundance between hippocampal subregions

To clarify the relationship between changing transcript levels and protein abundance, we next used DSP to profile select protein and phosphoprotein levels in each hippocampal subregion **(Table S3)**. A total of 87 proteins were quantified, using panels of antibodies targeting markers of neural and glial cell types, autophagy, cell death, MAPK and PI3K/AKT signaling pathways, and Alzheimer’s and Parkinson’s disease pathologies. Statistical comparisons between treatment groups and subregions mirrored those used for WTA. We first assessed the effects of SD itself (comparing SD vs. Sleep) with learning condition (CFC/HC) as a covariate. We observed increased levels of phosphorylated S6 (S235/S236) and phosphorylated ERK1/2 (42/44 MAPK; T202/Y204) across CA1, CA3, and DG hilus following SD, consistent with IEG expression changes we observed in these structures (**Fig. 3**). Together with our previous findings [17], these data suggest elevated activity in these subregions after SD. Additional SD-driven changes present in these regions include increased phosphorylated p90 RSK (T359/S363) and decreased phosphorylated Tau (T231) in CA1, increased phosphorylated Tau (S214) in both the hilus and CA3, and increased beta-secretase 1 (BACE1) and decreased myelin basic protein and S100B within CA3 only (**Fig. 8a; Table S3**). Somewhat surprisingly, no significant changes due to Sleep vs. SD were observed for proteins quantified in either of the DG blades when the learning condition was used as a covariate.

**Fig. 8:**
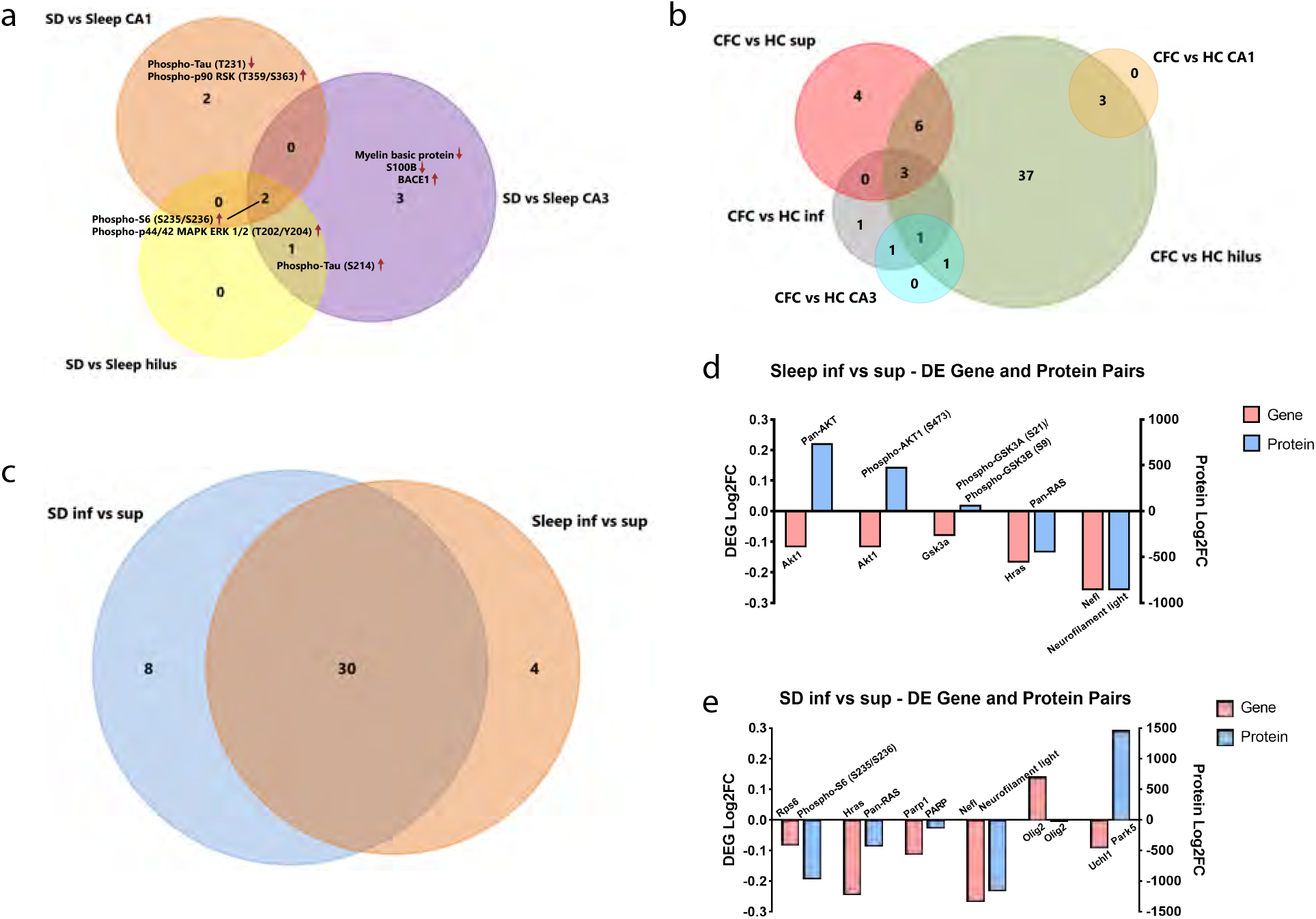
Learning and subsequent sleep or SD differentially affect protein expression in hippocampal subregions. **a,** Venn diagram of SD-altered proteins in CA1, hilus, and CA3. **b,** Venn diagram of CFC-altered proteins in hippocampal subregions. All fold change and FDR are reported in **Extended Data Table 3**. **c,** Venn diagram reflects the overlap (30 proteins) of inferior vs superior blade differentially expressed proteins under Sleep or SD condition. **d,** Log2FC of proteins differentially expressed between inferior and superior blade in the sleep mice and their corresponding DEGs for the same comparison in the WTA analysis. **e,** Log2FC of proteins differentially expressed between inferior and superior blade in the SD mice and their corresponding DEGs for the same comparison. All FDRs are reported in **Extended Data Table 4**.

We then compared protein expression between DG inferior and superior blades (Inferior vs. Superior), again using the learning condition (CFC/HC) as a covariate. 34 proteins differed between the blades in freely-sleeping mice; 38 were differentially expressed between the blades following SD (**Fig. 8c**; **Table S3**). 30 differentially-expressed proteins overlapped between SD and Sleep conditions (**Fig. 8c**). In freely-sleeping mice, 5 of the 34 differentially-expressed proteins had corresponding transcript-level differences by WTA, although the direction of expression differences between blades for transcript vs. protein did not always correspond. For example, pan-AKT and phosphorylated AKT1 (S473) levels were higher after sleep in the inferior blade, while *Akt1* transcript levels were higher in the superior blade. Phosphorylated GSK3A (S21) and GSK3B (S9) proteins were similarly expressed at higher levels in the inferior blade, while the *Gsk3a* transcript was more abundant in the superior blade. One plausible explanation for the differences in transcript vs. protein abundance is that these reflect differences in the rate of protein synthesis of transcripts between the blades. (**Fig. 8d; Table S4**).

In SD mice, 6 of the 38 proteins differentially expressed between the two DG blades also had transcript-level differences by WTA. Here again, the differences in protein and transcript levels occasionally moved in opposite directions. For example, Olig2 (a transcription factor that regulates oligodendrocyte, interneuron, and motor neuron development) protein was expressed at higher levels in the superior blade while the *Olig2* transcript was more abundant in the inferior blade. Park5 (ubiquitin C-terminal hydrolase L1) protein levels were higher in the inferior blade after SD, while corresponding transcript *Uchl1* was more abundant in the superior blade (**Fig. 8e**). As might be expected for SD mice (based on engram neuron reactivation and IEG expression levels; **Figs. 2** and **3**), phosphorylated S6 (S235/S236) and its corresponding transcript *Rps6* were both present at higher levels in the superior blade vs. inferior blade following SD. Similarly, pan-RAS GTPase, DNA repair enzyme poly-ADP-ribose polymerase (PARP), and neurofilament light, as well as corresponding transcripts *Hras, Parp1*, and *Nefl*, were all expressed at higher levels in the superior vs. inferior blade in SD mice. (**Fig. 8e**). Of these, pan-Ras and neurofilament light were consistently enriched in the superior blade regardless of the sleep state, indicating a plausible constitutive difference between the two blades (**Fig. 8d-e**; **Table S4**).

Lastly, we quantified changes in protein levels that were due to learning (comparing CFC vs. HC), using sleep condition (Sleep/SD) as a covariate (**Fig. 8b**). Three proteins were altered by learning in CA1 (neurofilament light, autophagy protein beclin 1, and MER tyrosine kinase) and in CA3 (Tau, phosphorylated Tau [T231], and apolipoprotein A-I). Learning altered slightly more proteins in the superior and inferior blades (13 and 6, respectively). Of these, 3 were significantly increased after learning in the inferior blade only (Tau, phosphorylated Tau [T199], phosphorylated Tau [T231]), and 10 were altered by learning in the superior blade only (including increases in ERK1/2, NeuN [a neuronal nuclear protein that regulates synaptic plasticity], MAP2, and Park5, and decreases in the amyloid-β regulating enzyme neprilysin, phosphorylated GSK3A [S21] and GSK3B [S9], phospho-AMPK-alpha [T172], phospho-PRAS40 [T246], p53, and beta-secretase 1). Surprisingly, the subregion with the most significantly-altered proteins after learning was the hilus, where (despite the lack of transcripts altered by CFC) 51 proteins were affected. 3 of these protein-level changes overlapped among all three DG subregions (superior, inferior blade and hilus) after CFC: MEK1 was significantly upregulated, while pan-RAS and phosphorylated ERK1/2 (42/44 MAPK; T202/Y204) were significantly downregulated.

Taken together, these data suggest that as is true for transcriptomic changes following learning and subsequent sleep or SD, protein abundance is also differentially affected by these processes within each dorsal hippocampal subregion. While the number of proteins and phosphoproteins profiled here was limited, these changes included effects on components of signaling pathways involved in activity-dependent plasticity, protein synthesis and ubiquitination, and responses to neuronal stress. These effects were also subregion-specific, with changes due to sleep vs. SD alone (regardless of prior learning) most prominent in CA1 and CA3 (and absent from the DG blades), and changes due to learning (regardless of subsequent sleep or SD) most prominent in the DG.

## Discussion

Our results demonstrate that post-learning DG engram neuron reactivation is state-dependent and occurs in a subregion-specific manner during post-learning sleep (**Figs. 1-2**). We also find that while most hippocampal subregions (DG hilus, CA1, and CA3) are more active during SD than during sleep, activation of DG inferior blade granule cells is selectively suppressed (**Fig. 3**). These subregion-specific findings inspired us to explore how the transcriptome within these regions is affected by single-trial CFC, and as a function of sleep vs. SD (**Figs. 4-7**). Our spatial transcriptomic analysis demonstrates that transcripts affected by learning and subsequent sleep state differ substantially between CA1, CA3, and DG subregions. We also find that when measured 6 h following CFC, learning-driven transcript changes are detectable only within the DG superior and inferior blade (despite persistent, and subregion-specific, changes in select proteins’ abundance in all regions). Together these findings present a picture of the hippocampus as a diverse landscape, where sleep loss has distinct effects in each region, and where following learning, persistent sleep-dependent, activity-driven gene expression changes are restricted to the DG.

This study adds to what is known about the function of hippocampal engram ensembles during memory encoding and consolidation, and more specifically how sleep contributes to this process. Engram neurons in DG are thought to be critical for contextual memory encoding [36] and recall (25, 26). Our data **(Fig. S1)** agree with recent findings that context-associated behaviors preferentially activate suprapyramidal blade DG granule cells [28, 29]. We find that context-activated neurons in the superior blade are preferentially reactivated during context re-exposure and same-context CFC **(Fig. 1d** and **j)**. Together, these findings suggest that context is more selectively encoded in DG by superior blade neurons than by inferior blade neurons.

Sleep-associated reactivation of engram neurons has been hypothesized as an important mechanism for memory consolidation (2, 3, 42). However, only a few recent studies have focused on the functional role of engram populations in the hours following learning (12, 28, 29). Here we find evidence for sleep-associated reactivation of DG engram neurons, with post-CFC sleep required for reactivation in the inferior blade (**Fig. 2e-f**). Because our data suggest that the inferior blade engram population is less selective for context than superior blade TRAP-labeled neurons, it is tempting to speculate that reactivation of inferior blade neurons during sleep provides mnemonic information about other aspects of the CFM engram – for example, the emotional salience of the context, rather than the context itself. This would be consistent with the phenotype observed after experimental post-CFC sleep disruption – i.e., a reduction in freezing behavior, suggesting a lack of fear association with the context (9, 11, 12) (**Fig. 2b**). Future studies will be needed to clarify the differential functional consequences of offline reactivation in superior vs. inferior blade engram populations. While technically challenging, this could theoretically be addressed by selectively activating or inhibiting one population or the other in the hours following CFC.

An unanswered question is *why* engram neuron populations in the two DG blades respond differently to post-learning sleep vs. SD. One possibility is that overall, granule cell activity levels differ between the blades during SD. Consistent with this, we find that Arc+ and Fos+ neuron numbers in DG inferior blade, but not the superior blade or hilus, consistently decrease after 6-h SD (**Fig. 2i**, **Fig. 3c-d**). Spatial transcriptomic profiling largely corroborates this – with activity-driven transcripts’ abundance reduced by SD in the inferior blade, and simultaneously increased in other hippocampal subregions (e.g., **Fig. 5b**, **d**, and **f**; **Fig. 6b**).

What neurobiological mechanisms underlie the selective SD-driven gating of inferior blade granule cells? Our present findings confirm constitutive differences in cell marker expression (e.g., higher *Npy* and *Penk* expression in superior blade vs. inferior blade; **Fig. 6b**), which are present between the two blades across conditions. Thus, there are apparent baseline differences in the cellular makeup of the two blades’ granule cell populations (32), which may impact the processes of learning and memory storage. The two blades are also known to differ with respect to their cortical inputs (43), with a greater number of lateral entorhinal inputs targeting the superior blade, and more numerous medial entorhinal inputs targeting the inferior blade. Thus, one possibility is that these excitatory presynaptic inputs are differentially active in the context of SD. Another important distinction is that the superior blade also has roughly half the interneuron density of the inferior blade (44). This latter finding suggests that selective inhibitory gating of the inferior blade could serve as a plausible mechanism by which engram neuron reactivation is suppressed during SD (20, 45).

We used spatial transcriptomics in this study as a tool to better understand the broader transcriptomic changes that are spatially and temporally associated with differences in overall activation, and engram neuron reactivation, of the two DG blades during SD. We find that, as was true across all 5 profiled hippocampal subregions (**Figs. 4** and **5; Fig. S5**), the two blades’ transcriptomic responses to SD were distinct (**Fig. 4**; **Figs. S5, S7,** and **S8**). Moreover, transcript differences between the blades differed substantially between Sleep and SD conditions (**Fig. 6**). Our analyses identified two key processes – regulation of actin cytoskeleton, and ribosomal biogenesis – which are significantly downregulated in the inferior blade after SD. Critically, these findings are consistent with available neuroanatomical and biochemical data, which have found disruptions in these pathways after SD, either across the hippocampus as a whole, or specifically within the DG (12, 15, 41, 46). Since both actin cytoskeletal remodeling and ribosomal biogenesis are critically associated with memory storage in the hippocampus (12, 15, 47, 48), disruption of both pathways in inferior blade at the transcriptional level could be sufficient to disrupt consolidation of CFM.

Our spatial profiling of dorsal hippocampus adds to a growing literature emphasizing the heterogeneity of SD effects of gene expression (12, 21) and neural activity (20, 21, 49, 50) in various brain structures. Collectively, these data demonstrate that sleep functions, particularly with regard to synaptic plasticity and cognition, cannot be assumed to be uniform across neuron (or other brain cell) types, or brain circuits (2, 51). It is worth noting that while the transcripts altered by SD varied between hippocampal subregions, with only partial overlap (**Figs. 4** and **5**), these SD-altered transcripts also only partially overlapped with transcripts altered by SD across whole hippocampus (18), and ribosome-associated transcripts isolated from specific hippocampal cell populations (12, 16) and subcellular fractions (12) after SD (**Figs. S12** and **S13**). This diversity of SD effects is also present, in our hands, between subregions at the level of protein expression (**Fig. 8a**). Given this diversity – with SD effects on transcripts and proteins varying with cell compartment, type, and structure, it is plausible that our present findings could by biased due to technical aspects of our spatial profiling strategy. For example, because only the cell body layers are profiled in DG, CA3, and CA1, it is likely that transcripts which are efficiently transported out of the soma (i.e., into dendrites and axons) are undersampled using our technique (12, 35)(52, 53). In addition, we cannot discriminate between transcript changes occurring within the principal (i.e., pyramidal or granule) cell bodies, those occurring in presynaptic terminals of other excitatory, inhibitory, or neuromodulatory neurons which terminate in those layers, and those occurring in microglia, astrocytes or oligodendrocytes. For this reason, additional experiments will be needed to precisely identify the cellular sources for some of transcript and protein changes we report here after SD – e.g. differential expression of Olig2 transcript and protein between the two DG blades, which could be driven by changes in mature oligodendrocytes, precursor cells, or neurons. Another example where the cellular source of SD-driven transcript changes is unclear is within the superior blade, where transcripts encoding synaptic release regulatory machinery were downregulated after SD. While this may be due to transcript changes in the granule cells themselves, it is also plausible that this effect of SD is due to altered transcripts in presynaptic terminals located within the superior blade.

We also used spatial profiling as a strategy to identify subregion-specific changes associated with storage of a new memory (CFM) in the hippocampus. One surprising finding from this analysis was the relative paucity of detectable learning-induced transcript changes in the hippocampus at 6 h post-CFC, and that these were present only within the DG blades (**Fig. 7**). This may be due in part to the fact that measurements occurred several hours after the single-trial learning event; it is plausible that many transcriptional responses to memory encoding are relatively transient, and for that reason are no longer detectable by this 6-h time point. While additional studies will be needed to clarify this point, what is certain is that sustained transcriptomic responses to CFC are restricted to the two DG blades. Critically, we find that transcripts that remain altered several hours after CFC include those critically involved in glutamatergic and GABAergic synaptic functions – which are generally upregulated and downregulated, respectively, in the superior blade after CFC (**Fig. 7; Fig. S11**). Identified CFC-driven transcript changes in inferior blade, while fewer, followed a similar overall pattern – i.e., transcripts involved in glutamatergic synapse biogenesis were upregulated, while GABA receptor transcripts were downregulated, after CFC. This strongly supports the hypothesis that excitatory-inhibitory balance changes in specific brain circuits accompanies, and benefits, long-term memory storage (54)(20, 55, 56). The finding that these changes are constrained to DG, among all subregions profiled, is consistent with the prior finding of sustained transcriptomic changes occurring specifically within DG engram neurons 24 h post-CFC (57), and may reflect unique dynamics of the DG network (or the engram neurons within it), which could plausibly outlast changes after learning in other hippocampal structures (11, 58, 59). This interpretation could reconcile the findings that while CFC-associated transcriptomic changes were absent in CA3 and CA1 at the 6-h timepoint, while protein abundance changes due to CFC were still detected (**Fig. 8**).

Together, our data advance the findings of previous studies (20, 21, 31, 32, 60, 61) highlighting the heterogeneity of hippocampal subregions’ function in the context of learning, and response to subsequent sleep or sleep loss. The present findings suggest that the influences of sleep and SD on hippocampally-mediated memory consolidation are linked to subregion-specific changes in both engram neuron reactivation and biosynthetic events related to protein synthesis regulation, synaptic signaling, and remodeling of neuronal cytoskeletal structures.

## Materials and Methods

### Animal handling and husbandry

All animal husbandry and experimental procedures were approved by the University of Michigan Institutional Animal Care and Use Committee. Mice from 3 to 6 months old were used for all experiments. With the exception of a 3-day period of constant dark housing following 4- hydroxytomaxifen (4-OHT) administration (see below), mice were maintained on a 12 h: 12 h light/dark cycle (lights on at 9AM) with *ad lib* food and water throughout the study. cfos-CRE^ER^ mice (31) B6.129(Cg)-Fos*^tm1.1(cre/ERT2)Luo^*/J; Jackson) were crossed to B6.Cg-Gt(ROSA)26Sor*^tm9(CAG-tdTomato)Hze^*/J (Jackson) mice to induce CRE recombinase-mediated expression of tdTomato (*cfos::tdTomato*). Mice were individually housed with beneficial environmental enrichment in standard caging 4 days prior to genetic tagging of engram cells and/or contextual fear conditioning (CFC), and were habituated to daily handling (2-5 min/day) for 3 days prior to the experiments.

### Genetic labeling of hippocampal engram cells

On the day of genetic labeling, starting at lights on (ZT0), mice were individually placed in one of the two novel contexts (Context A or Context B). Context A was a 24 × 24 × 23 cm square arena with metal grid floor, scented with an all-purpose sponge soaked with 5 ml Lysol (lemon breeze scent) attached to the lid of the chamber. Context A was surrounded by 4 LED monitors presenting a 135° flickering oriented grating stimulus (100% contrast, 0.05 cycles/deg, flickering at 1 Hz). Context B was a 23 × 23 × 22 cm cylindrical arena with a pink glossy plastic floor scented with an all-purpose sponge soaked with 1 ml 1% ethyl acetate attached to the lid of the chamber. Context B was surrounded by 4 LED monitors presenting either a vertical oriented grating stimulus or a dark screen. For genetic labeling, immediately following 15 min of free novel context exploration, mice received an i.p. injection of 4-OHT (50 mg/kg in corn oil). They were then returned to their home cage, which was placed in complete darkness inside a sound-attenuated chamber over the next 3 days to minimize non-specific TRAP-based neuronal labeling (62). 3 days following 4-OHT administration, mice were returned to a normal 12 h: 12 h LD cycle for an additional 3 days prior to behavioral experiments.

### Behavioral procedures

#### For CFC behavioral experiments

(**Fig. 2a-b**), At ZT0, male and female C57BL/6J mice (Jackson) or WT siblings of *cfos::tdTomato* mice underwent single-trial CFC as described previously (11, 58, 59). Briefly, mice were placed in Context A and were allowed 3.5 min of free exploration time prior to delivery of a 2-s, 0.75 mA foot shock through the chamber’s grid floor. After 5 min total in the chamber, mice were returned to their home cage, where they were either allowed *ad lib* sleep or were sleep deprived (SD) under normal room light for the first 6 h following training, using the gentle handling procedures (including cage tapping, nest material disturbance, and light touch with a cotton-tipped applicator when necessary). After SD, all mice were allowed *ad lib* sleep. 24 h following CFC, at lights on (ZT0; next day) mice were returned to Context A for 5 min to assess CFM. CFM was measured quantitatively as % time spent freezing during re-exposure to Context A using previously established criteria (63) (crouched, rigid posture with no head or whisker movement). Two scorers blinded to behavioral conditions independently quantified periods of freezing behavior prior to shock during CFC and during CFM testing.

#### For TRAP labeling and context re-exposure experiments

(**Fig. 1a-f**), at ZT0 on the day of re-exposure, male and female *cfos::tdTomato* mice that underwent TRAP labeling 6 days previously (following Context A exploration) were either returned to Context A, or were placed in distinct Context B. Following 15 min of free exploration, all mice were returned to their home cage inside of a dark, sound-attenuated chamber to minimize interference and disturbances. 90 min after the second context exposure, mice were sacrificed via an i.p. Injection of Euthasol and were transcardially perfused with ice-cold 1× PBS followed by 4% paraformaldehyde.

#### For context re-exposure with CFC experiments

(**Fig. 1g-l**), at ZT0 on the day of CFC, male *cfos::tdTomato* mice that underwent TRAP labeling 6 days previously (following either Context A or Context B exploration) underwent single-trial CFC in Context A as described above, after which they were returned to their home cage in a dark, sound-attenuated chamber. 90 min after CFC, mice were sacrificed and perfused as described above.

#### For context re-exposure with CFC followed by Sleep/SD

(**Fig. 2c-i**), at ZT0 on the day of the experiment, male *cfos::tdTomato* that underwent TRAP labeling 6 days previously (following Context A exploration) underwent single-trial CFC in Context A as described above. All mice were then returned to their home cage, and either were allowed *ad lib* sleep or were sleep deprived (SD) under normal room light for the next 6 h using gentle handling procedures (including cage tapping, nest material disturbance, and light touch with a cotton-tipped applicator when necessary). Immediately following the 6-h post-CFC sleep or SD window, mice were sacrificed and perfused as described above.

For spatial profiling experiments in **Figs. 3-6**, at ZT0 on the day of the experiment, male C57BL/6J mice (Jackson) underwent single-trial CFC in Context A, then were returned to their home cage and either were allowed *ad lib* sleep or were sleep deprived (SD) under normal room light for 6 h as described above. Immediately after SD, all mice were sacrificed and perfused as described above.

### Immunohistochemistry

Immediately following perfusions, brains were dissected and post-fixed at 4°C in 4% paraformaldehyde (PFA) for 24 hours. Post-fixed brains were then sectioned coronally at 80-100 μm using a vibratome (Leica VT1200 S). Sections containing dorsal hippocampus were blocked in PBS with 1% TritonX-100 and 5% normal donkey serum overnight, then incubated at 4°C for 3 days in rabbit-anti-cfos 1:1000 (Abcam; ab190289) and either goat-anti-tdTomato 1:600 (SICGEN; AB8181-200) or guinea pig-anti-Arc 1:500 (Synaptic Systems; 156004). Sections were then incubated with secondary antibodies at 4°C for 2 days in CF™ 633 Anti-Rabbit IgG H+L 1:1000 (Sigma-Aldrich; SAB4600132), Alexa Fluor® 488 AffiniPure Donkey Anti-Goat IgG H+L 1:800 (Jackson ImmunoResearch; 705-545-003), DAPI (Sigma-Aldrich D9542) or CF™555 Anti-Guinea Pig IgG H+L 1:1000 (Sigma-Aldrich; SAB4600298). Immunostained sections were coverslipped in ProLong Gold antifade reagent (ThermoFisher; P36930) and were imaged using a Leica SP5 upright laser scanning confocal microscope.

### Image quantification

Images of immunostained hippocampi were obtained as 20× z-stacks (step size = 5 μm) spanning the thickness of each brain slice. Settings were fixed for each imaging session. For data reported in **Fig. 1**, 3-6 dorsal hippocampal DG sections were quantified from each mouse. For data reported in **Fig. 2**, 7 DG sections were quantified per mouse. For data reported in **Fig. 3**, 5 sections of DG, and 3 sections of both CA1 and CA3 were quantified per mouse. Fluorescence images were analyzed using Fiji (64). For CA1, pyramidal layer Arc and cFos immunolabeling was quantified as average pyramidal layer fluorescence intensity minus average background intensity. In each image, the entire CA1 pyramidal cell body layer as one region of interest (ROI), and background fluorescence ROIs were outlined in adjacent regions with autofluorescence but without cFos and Arc immunolabeling; mean intensity of each ROI was obtained using Fiji. For DG and CA3 cFos+ and Arc+ cell quantification, maximum intensity z-projection was applied to each image stack, followed by adjusting the threshold to identify pixels above 1% of the intensity distribution; this yielded a binary image without background autofluorescence. Cell counting was then performed by a scorer blinded to experimental condition using Fiji. The overlap between tdTomato and cFos was quantified using cFos channel-thresholded consecutive single plane images to verify overlap between cFos and tdTomato signals within neuronal cell bodies.

### Statistical analysis

Statistical analyses in **Figs. 1, 2**, and **3** were performed using GraphPad Prism (version 9). Mann Whitney tests were used for analysis of cFos+, tdTomato+ neurons, or overlap percentages between pairs of treatment groups (**Fig. 1c-f**, **Fig. 1i-l**, **Fig. 2e-i, and Fig. 3**). Wilcoxon matched-pairs signed rank tests were used for analysis of overlap percentages in different subregions within the same animals (**Fig. 2f** and **h**; superior vs. inferior). A Student’s t-test was used for behavioral analysis of CFM (**Fig. 2b**). For each specific comparison, the statistical tests used are listed in the figure legend. Within figures, *, **, ***, and **** indicate *p* < 0.05, *p* ≤ 0.01, *p* ≤ 0.001, *p* ≤ 0.0001, respectively.

### GeoMx Digital Spatial Profiling (DSP) slide preparation

A total of 16 PFA-fixed brains (from *n* = 4 male mice for each condition: HC-Sleep, HC-SD, CFC- Sleep, and CFC-SD) were cryosectioned at 10 μm thickness less than 2 weeks prior to GeoMx Mouse Whole Transcriptome Assay (WTA) and protein panels. Four brain sections containing dorsal hippocampus (1 from each experimental condition) were placed onto each slide (Fisherbrand Superfrost Plus) to reduce technical artifacts introduced on individual slides during the slide preparation process. Slides were stored in −80°C until being prepared for WTA. DSP was performed as described in detail in Merritt et al. (2020) (65). Briefly, slides with fixed frozen brain sections (1 section per mouse) were baked at 60°C for 1 h, then were post-fixed with 4% PFA in 1× PBS at 4°C for 15 min. Dehydration was performed for 5 min in 50% EtOH, 2 × 5 min in 70% EtOH and 5 min in 100% EtOH, followed by antigen retrieval using boiling 10 mM sodium citrate, pH 6.0 (Sigma Aldrich C9999) for 5 min. Protease III (ACD Cat#322337) was added to brain sections at 40°C for 30 min to expose RNA targets, then *in situ* hybridizations with mouse WTA panel (20175 targets) were performed in Buffer R (NanoString); slides were covered by HybriSlips (Grace Biolabs) and incubated at 37°C overnight. The following day, two 25-min stringent washes were performed using 50% formamide and 2 × SSC, followed by an additional two 2-min washes using 2 × SSC. Brain sections were then blocked with Buffer W (NanoString) for 30 min at room temperature. SYTO 13 (NanoString, 1:100 in Buffer W) was added to each slide for 1 h in a humidified chamber at room temperature; this fluorescent nuclear stain was used to identify borders of hippocampal subregions. Slides were then briefly washed twice using 2 × SSC and were immediately loaded on the GeoMx Digital Spatial Profiler.

For protein panels, slides were baked at 60°C for 1 h followed by dehydration and antigen retrieval. The slides were then blocked using NanoString blocking buffer W for 1 h. A NanoString protein antibody cocktail including Mouse Protein Core + Controls, Mouse PI3K/AKT Module, Mouse MAPK Signaling Module, Mouse Cell Death Module, Mouse Neural Cell Typing Module, Mouse AD Pathology Module, Mouse AD Pathology Extended Module, Mouse PD Pathology Module, Mouse Glial Subtyping Module, and Mouse Autophagy Module (NanoString Technologies) was added to the sections followed by overnight incubation at 4°C. The next day, SYTO 13 was applied to each slide before loading slides on the GeoMx Digital Spatial Profiler. During slide washes for the protein panel experiment, one HC-sleep, one HC-SD, and one CFC- Sleep brain section were lost, and were excluded from subsequent experiments. Another HC-SD brain section provided only one CA1 subregion that was usable for both WTA and protein experiments.

Each slide was imaged using a 20 × objective, and bilateral ROIs from each hippocampal subregion were selected using SYTO 13 staining using the GeoMx software. After ROI selection, UV light was applied to each ROI to achieve subregion-specific indexing oligonucleotide cleavage. The released oligonucleotides were collected via a microcapillary and dispensed into a 96-well plate; these were dried at room temperature overnight. Sequencing libraries were then prepared per the manufacturer’s protocol. Libraries were sequenced on an Illumina NovaSeq 6000 with an average depth of 5.3 million raw reads per ROI.

### GeoMx data analysis

Raw FASTQ files were processed using Nanostring’s Automated Data Processing Pipeline. This pipeline includes adapter trimming, aligning stitched paired-end reads to barcodes in the reference, and removing PCR duplicates based on the Unique Molecular Identifier (UMI) in each read (GeoMx DSP NGS Readout User Manual, https://university.nanostring.com/geomx-dsp-ngs-readout-user-manual/1193408). All QC and downstream differential expression analyses were performed in R version 4.2.0 (2022-04-22) using the NanoString-developed GeoMxTools package (versions 1.99.4 for WTA and 3.0.1 for protein) (66). Plots were generated using either built-in GeoMxTool functions or the ggplot2 package (version 3.3.6). For the mouse WTA panel (v1.0), each gene target is mapped to a single probe with a total of 20,175 probes, 210 of which are negative probes that target sequences not present in the transcriptome. For each ROI, a Limit of Quantification (LOQ; the geometric mean of negative probes × the geometric standard deviation of negative probes raised to a power of 2) was calculated. All 128 selected ROIs exceeded a 15% gene detection rate. Gene targets that failed to be detected above the LOQ in at least 10% of the segments were removed as recommended, leaving 11508 gene targets in the final filtered WTA dataset.

For mouse protein panels (**Table S3**), 119 out of the initial 121 ROIs passed Nanostring recommended QC thresholds and were used for DE analyses. In total, 93 protein targets were tested, 3 of which are negative controls. Signal to background ratio for each protein target was calculated; 6 protein targets were removed due to low detection levels, leaving 87 protein targets in the final filtered protein data.

Filtered WTA and protein data were normalized using the third quartile normalization method. Linear mixed-effect models (LMMs) were used to perform differential expression analysis. Depending on the comparison, combinations of fixed effects of sleep condition (Sleep/SD), learning condition (CFC/HC), and random effects of slide (taking into account variations during slide processing steps) were adjusted for. Genes, transcripts, and protein targets that have an FDR < 0.1 are considered differentially expressed (DE). Venn diagrams of overlapping DEGs/DE proteins between regions/conditions were made using FunRich (67). Gene ontology (GO), KEGG pathway, and predicted upstream regulator analyses were performed using Advaita Bio’s iPathwayGuide (https://advaitabio.com/ipathwayguide), using an FDR < 0.1 multiple comparisons *p* value correction for all analyses. A smallest common denominator pruning method (iPathwayGuide) was used to identify GO terms, due to the high number of DEGs and overlapping GO terms identified. FDR-based p-value correction was used for all other analyses.

## Data and code availability

Processed sequencing data and raw data for both RNA and protein are available in the Gene Expression Omnibus (GEO, https://www.ncbi.nlm.nih.gov/geo/) under accession number GSE229082. Code for all data analyses and input files are available on GitHub for download, at: https://github.com/umich-brcf-bioinf-projects/Aton_saton_CU8-GeoMx.

## Acknowledgements

Figures illustrating behavioral paradigms were created with BioRender. We are grateful to the staff of the Advanced Genomics Core at the University of Michigan for assistance with GeoMx slide preparation, library preparation and next-generation sequencing.

## Supplemental Figure Legends

**Fig. S1. a,** Numbers of tdTomato+ and (**b**) cFos+ neurons in the DG granule cell layer were similar in A to A and A to B mice. Values indicate mean ± s.d.. **c,** tdTomato+ neuron densities in A to A and A to B mice were similar for comparisons within the same blade, e.g. superior A to A vs. superior A to B. For both A to A and A to B scenarios, the inferior blade had fewer tdTomato+ neurons compared the superior blade (** ** indicates *p* = 0.0039 and *p* = 0.0039, respectively, Wilcoxon matched-pairs signed rank test). **d,** cFos+ neuron densities in A to A and A to B scenarios showed a similar pattern, with the inferior blade having fewer cFos+ neurons than the superior blade (** indicates *p* = 0.0039 and *p* = 0.0078, respectively, Wilcoxon matched-pairs signed rank test). **e,** Numbers of tdTomato+ and **f**, cFos+ neurons in the DG granule cell layer were similar in A to A-CFC and A to B-CFC mice. Values indicate mean ± s.d.. **g-h,** tdTomato+ and cFos+ neuron densities in the superior and inferior blades were comparable in A to A-CFC and A to B-CFC mice. Again, the inferior had fewer tdTomato+ neurons (* indicates *p* = 0.0156 and *p* = 0.0156, respectively, Wilcoxon matched-pairs signed rank test) and fewer cFos+ neurons compared the superior blade (* indicates *p* = 0.0156 and *p* = 0.0156, respectively, Wilcoxon matched-pairs signed rank test), in both scenarios. **i,** Numbers of tdTomato+ and **j**, cFos+ neurons in the DG granule cell layer were similar in Sleep and SD mice. **k**, tdTomato+ neuron densities in the superior blade or inferior blade were comparable in Sleep and SD mice. The inferior blade had fewer tdTomato+ neurons compared the superior blade under both Sleep and SD conditions (** indicates *p* = 0.0039 and ** *p* = 0.0020, respectively, Wilcoxon matched-pairs signed rank test). Values indicate mean ± s.d..

**Fig. S2. a,** Representative image of TRAPed hilar tdTomato+ neurons. Neuronal morphology for TRAPed hilar neurons was distinct from labeled granule cells. **b,** Percentage of cFos+/tdTomato+ cells in the DG hilus, as well as the numbers of (**c**) tdTomato+ and (**d**) cFos+ neurons, were similar between A to A and A to B mice. **e,** Percentage of cFos+/ tdTomato+ cells in the DG hilus, as well as the numbers of (**f**) tdTomato+ and (**g**) cFos+ neurons, were similar in A to A-CFC and A to B- CFC mice. **h,** Percentage of cFos+/tdTomato+ cells in the DG hilus, as well as the numbers of (**i**) tdTomato+ neurons, were similar in Sleep and SD mice. Values indicate mean ± s.d..

**Fig. S3. a,** Representative images showing overlap of tdTomato (***magenta***) and cFos protein (***green***) in the pyramidal layer of CA1 and **b,** CA3. Scale bar = 100 μm. **c,** Percentage of cFos+/tdTomato+ cells in the CA1 pyramidal layer was significantly higher following SD (*n* = 9) than sleep (n = 9). * indicates *p* = 0.0367, Mann Whitney test. **d,** Numbers of tdTomato+ neurons in the CA1 pyramidal layer were similar in sleep and SD mice. **e,** Percentage of cFos+/tdTomato+ cells in the CA3 pyramidal layer, as well as **f,** the numbers of tdTomato+ neurons, were similar in Sleep and SD mice. Values indicate mean ± s.d..

**Fig. S4. a,** Principal component analysis of all transcripts showing regions of interest (ROIs) strongly stratified according to hippocampal subregion, regardless of treatment group. **b,** Nuclei count in hippocampal subregions as number of nuclei per region of interest (ROI) from GeoMx

**Fig. S5 a,** UpSet plot and **b,** Venn diagram of overlapping SD-altered DEGs between each of the 4 principal cell body subregions (CA1, CA3, Inferior blade, Superior blade). The number of total DEGs for each subregion is shown on the right side of the UpSet plot. Dots with connecting lines indicate overlap of DEGs between one or more subregions, and the corresponding overlap can be found in the Venn diagram. Individual dots with no connecting lines indicate DEGs only affected in the specific subregion. The number of DEGs for a specific overlap is shown on the top of the UpSet plot. **c,** UpSet plot of only SD-upregulated DEGs between each hippocampal subregion. **d,** UpSet plot of only SD-downregulated DEGs between each hippocampal subregion. **e,** Heatmap of the 53 DEGs altered by SD across CA1, CA3, DG superior and inferior blades.

**Fig. S6. a,** Gene ontology cellular component term, ER chaperone complex (GO:0034663), was significantly impacted in the DG superior, inferior blade, hilus, and CA3 following SD. **b,** Expression level of SD vs Sleep DEGs annotated to GO:0034663 in superior blade, inferior blade, hilus, CA1, and CA3.

**Fig. S7. a,** SD altered DEGs annotated to Glutamatergic synapse and Ribosome pathways in the superior blade of DG. **b,** SD altered DEGs annotated to Retrograde endocannabinoid signaling and Oxytocin signaling pathways in the inferior blade of DG. **c,** SD altered DEGs annotated to Serotonergic synapse and GABAergic synapse in the CA1.

**Fig. S8.** Diagrams of SD vs. sleep impacted the Circadian entrainment KEGG pathway (KEGG: 04713) in the superior blade, inferior blade, and CA1. The perturbation accounts both for the gene’s measured fold change, and for the computed accumulated perturbation propagated from fold changes to any significantly altered upstream regulator genes, with the highest negative perturbation shown in dark blue, and the highest positive perturbation shown in dark red. The legend describes the values on the gradient for each subregion.

**Fig. S9.** SD vs. Sleep DEG overlap between hippocampal subregions. **a**, Illustration of the comparison and number of DEGs for SD vs Sleep in each hippocampal subregion. **b,** 9 DEGs overlapped between CA1 and hilus, all of them were regulated in the same direction (i.e., were similarly up or downregulated) by SD. **c,** 12 DEGs overlapped between superior blade and hilus; all were regulated in the same direction by SD. **d,** 7 DEGs overlapped between CA3 and hilus; all were regulated in the same direction by SD. **e**, 960 DEGs overlapped between superior and inferior blades; all were regulated in the same direction by SD. **f,** 134 DEGs overlapped between CA1 and CA3; all were regulated in the same direction by SD.

**Fig. S10. a,** Inferior vs. Superior blade KEGG pathways under SD condition. **b,** Inferior vs. Superior blade differentially expressed pathway genes in the Regulation of actin cytoskeleton pathway.

**Fig. S11. a,** Illustration of the CFC vs HC comparison. **b,** Significant GO terms mapped for the DEGs altered by CFC in the superior blade. **c,** CFC downregulated DEGs annotated to synaptic membrane. **d,** CFC upregulated DEGs annotated to synaptic membrane.

**Fig. S12**. Venn diagrams show the overlap for transcripts altered by SD in each hippocampal subregion and those previously reported in **a**, RNAseq of whole hippocampus following SD (Gaine et al., 2021), and **b**, translating ribosome affinity purification-seq from Camk2a+ neurons following SD (Lyons et al., 2020).

**Fig. S13**. Overlap of SD vs Sleep DEGs with previously-characterized SD-altered translating ribosome affinity purification transcripts. Venn diagrams indicate the overlap for transcripts altered by SD in five different hippocampal subregions in the present study, compared with our previous translating ribosome affinity purification-seq study, where DEGs were identified from whole hippocampus, subsampling **a**, Camk2a+ neurons, **b**, highly active, pS6+ neurons, and **c**, input (without subsampling) following SD (Delorme et al., 2021), for both cytosolic and membrane- associated fractions.

