## Supplemental material for "Sleep-dependent engram reactivation during hippocampal memory consolidation is associated with subregion-specific biosynthetic changes"

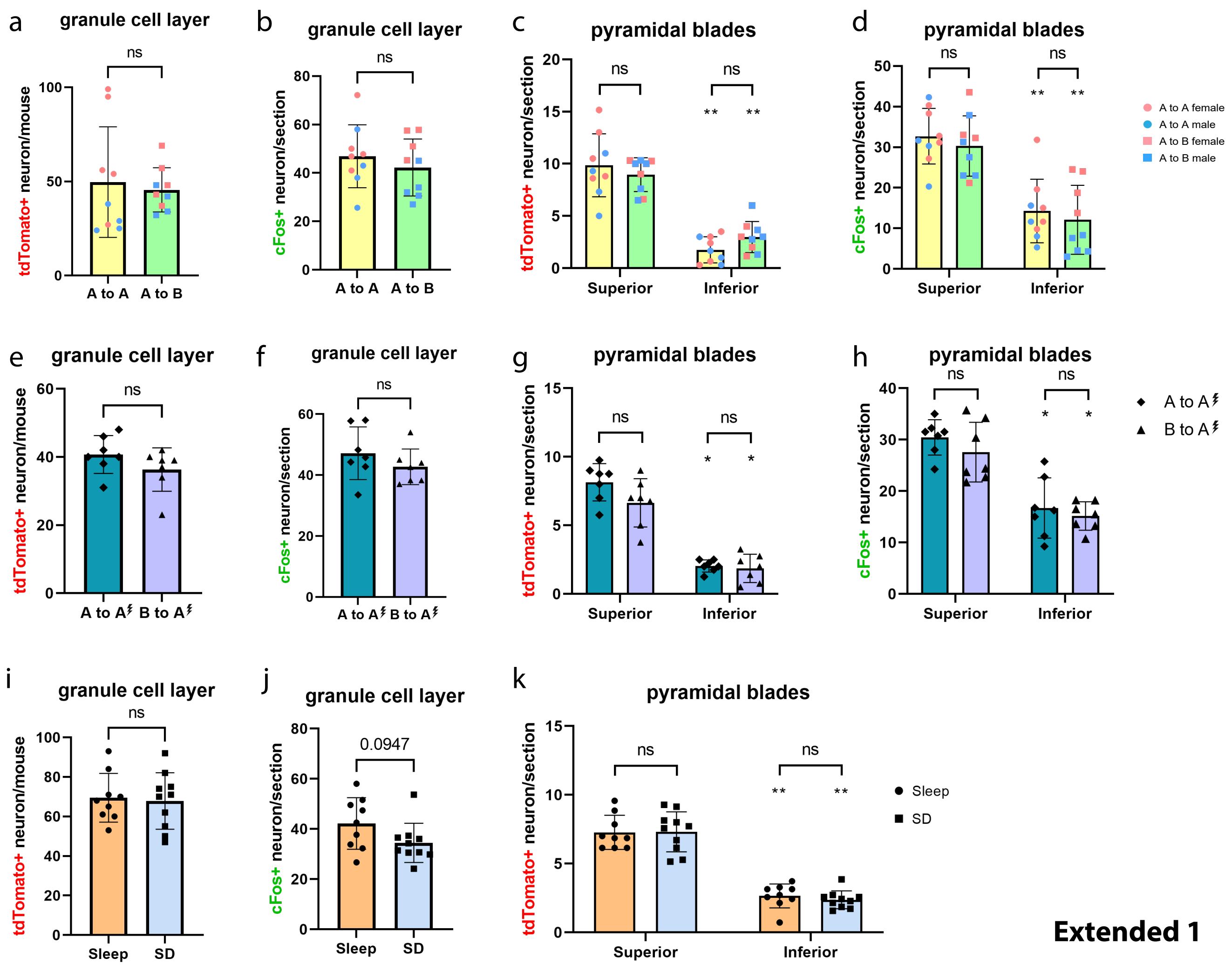

a
granule cell layer
b
granule cell layer
c
pyramidal blades
d
pyramidal blades
A to A female
A to A male
A to B female
A to B male
e
granule cell layer
f
granule cell layer
g
pyramidal blades
h
pyramidal blades
i
granule cell layer
j
granule cell layer
k
pyramidal blades
Sleep
SD
tdTomato+ neuron/mouse
cFos+ neuron/section
tdTomato+ neuron/section
Superior
Inferior
A to A
A to B
ns
0.0947


Extended 1

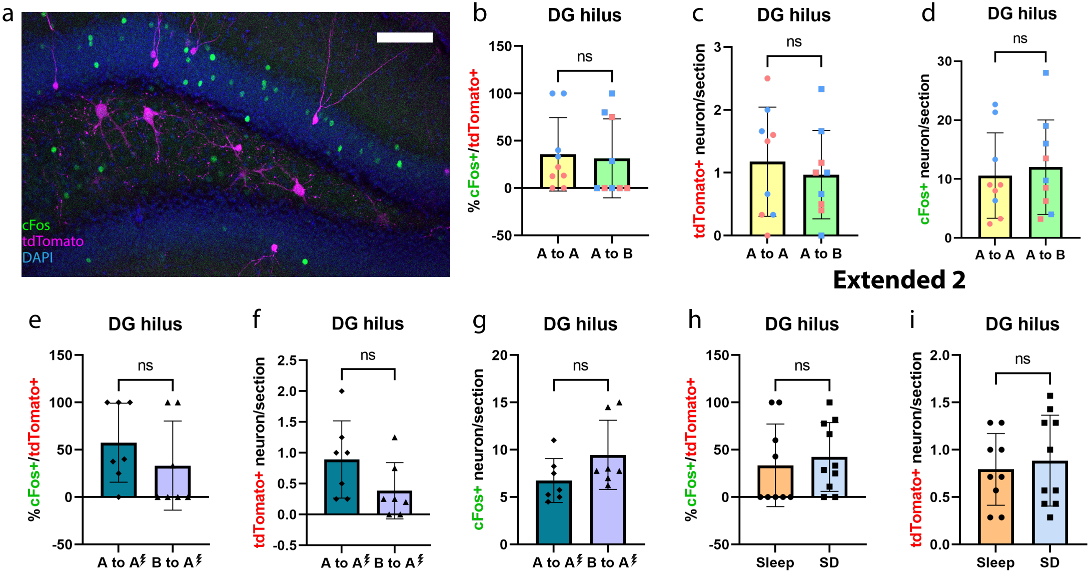
a
cFos
tdTomato
DAPI
b
DG hilus
% cFos+/tdTomato+
150
100
50
0
-50
ns
A to A
A to B
c
DG hilus
tdTomato+ neuron/section
3
2
1
0
ns
A to A
A to B
d
DG hilus
cFos+ neuron/section
30
20
10
0
ns
A to A
A to B
e
DG hilus
% cFos+/tdTomato+
150
100
50
0
-50
ns
A to A
B to A
f
DG hilus
tdTomato+ neuron/section
2.5
2.0
1.5
1.0
0.5
0.0
-0.5
ns
A to A
B to A
g
DG hilus
cFos+ neuron/section
20
15
10
5
0
ns
A to A
B to A
h
DG hilus
% cFos+/tdTomato+
150
100
50
0
-50
ns
Sleep
SD
i
DG hilus
tdTomato+ neuron/section
2.0
1.5
1.0
0.5
0.0
ns
Sleep
SD

**Extended 2**

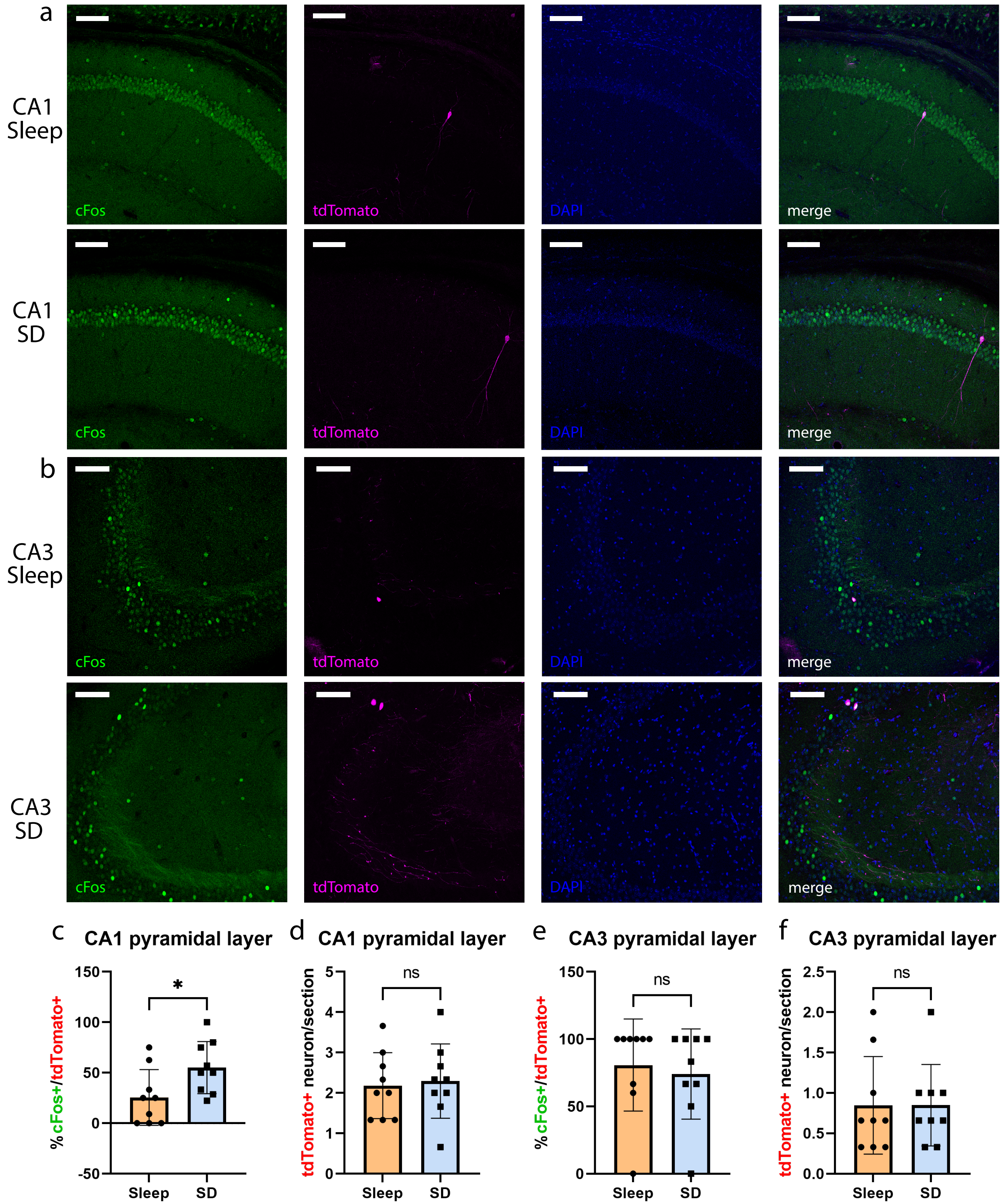
a
CA1
Sleep
cFos
tdTomato
DAPI
merge
CA1
SD
cFos
tdTomato
DAPI
merge
b
CA3
Sleep
cFos
tdTomato
DAPI
merge
CA3
SD
cFos
tdTomato
DAPI
merge
c
CA1 pyramidal layer
% cFos+/tdTomato+
150
100
50
0
-50
*
Sleep
SD
d
CA1 pyramidal layer
tdTomato+ neuron/section
5
4
3
2
1
0
ns
Sleep
SD
e
CA3 pyramidal layer
% cFos+/tdTomato+
150
100
50
0
ns
Sleep
SD
f
CA3 pyramidal layer
tdTomato+ neuron/section
2.5
2.0
1.5
1.0
0.5
0.0
ns
Sleep
SD

**Extended 3**

### Extended 4

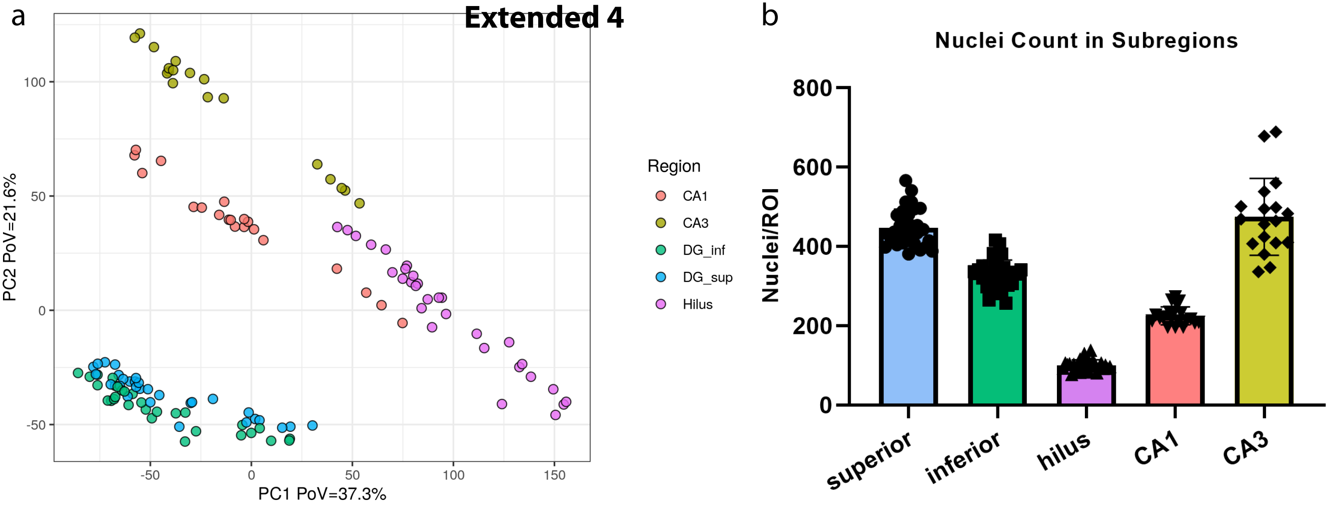
a
PC2 PoV=21.6%
PC1 PoV=37.3%
Region
CA1
CA3
DG_inf
DG_sup
Hilus
b
Nuclei Count in Subregions
Nuclei/ROI
superior
inferior
hilus
CA1
CA3

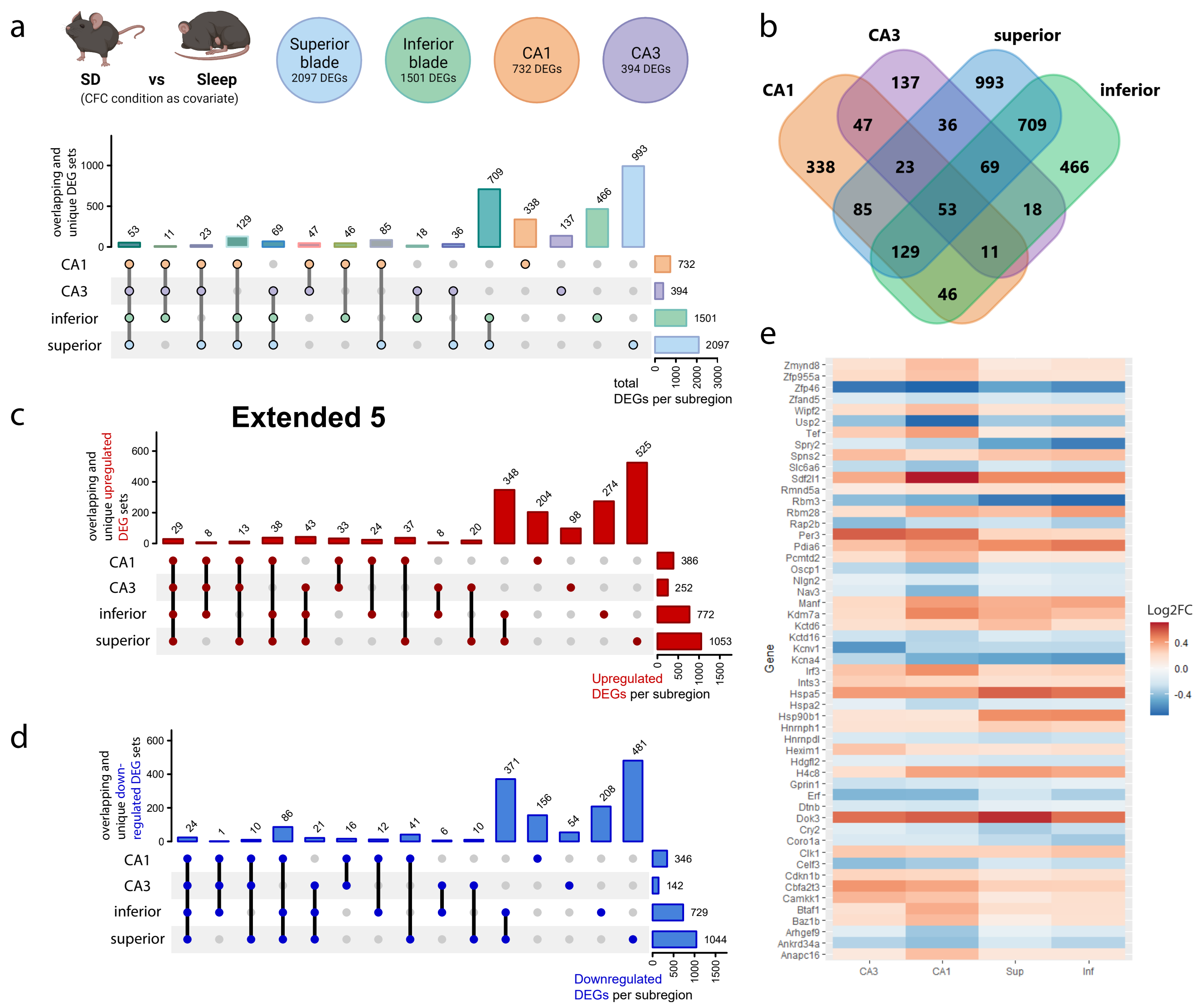
a
SD
vs
Sleep
(CFC condition as covariate)
Superior blade 2097 DEGs
Inferior blade 1501 DEGs
CA1 732 DEGs
CA3 394 DEGs
overlapping and unique DEG sets
53 11 23 129 69 47 46 85 18 36 709 338 137 466 993
CA1
CA3
inferior
superior
732
394
1501
2097
total DEGs per subregion
b
CA1
CA3
superior
inferior
137
993
47
36
709
338
23
69
466
85
53
18
129
11
46
c
Extended 5
overlapping and unique upregulated DEG sets
29 8 13 38 43 33 24 37 8 20 348 204 98 274 525
386
252
772
1053
Upregulated DEGs per subregion
d
overlapping and unique down-regulated DEG sets
24 1 10 86 21 16 12 41 6 10 371 156 54 208 481
346
142
729
1044
Downregulated DEGs per subregion
e
Gene
Zmynd8
Zfp955a
Zfp46
Zfand5
Wipf2
Usp2
Tef
Spry2
Spns2
Slc6a6
Sdf2l1
Rmnd5a
Rbm3
Rbm28
Rap2b
Per3
Pdia6
Pcmtd2
Oscp1
Nlgn2
Nav3
Manf
Kdm7a
Kctd6
Kctd16
Kcnv1
Kcna4
Irf3
Ints3
Hspa5
Hspa2
Hsp90b1
Hnrnph1
Hnrnpdl
Hexim1
Hdgfl2
H4c8
Gprin1
Erf
Dtnb
Dok3
Cry2
Coro1a
Clk1
Celf3
Cdkn1b
Cbfa2t3
Camkk1
Btaf1
Baz1b
Arhgef9
Ankrd34a
Anapc16
CA3
CA1
Sup
Inf
Log2FC
0.4
0.0
-0.4

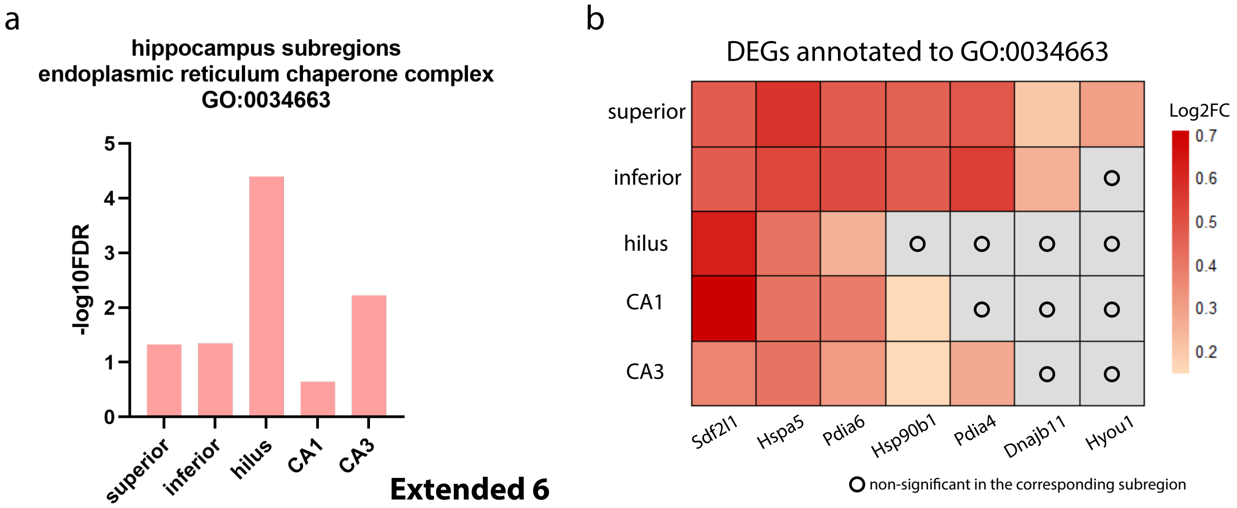
a
hippocampus subregions
endoplasmic reticulum chaperone complex
GO:0034663
-log10FDR
0
1
2
3
4
5
superior
inferior
hilus
CA1
CA3
b
DEGs annotated to GO:0034663
superior
inferior
hilus
CA1
CA3
Sdf2l1
Hspa5
Pdia6
Hsp90b1
Pdia4
Dnajb11
Hyou1
Log2FC
0.7
0.6
0.5
0.4
0.3
0.2
non-significant in the corresponding subregion

**Extended 6**

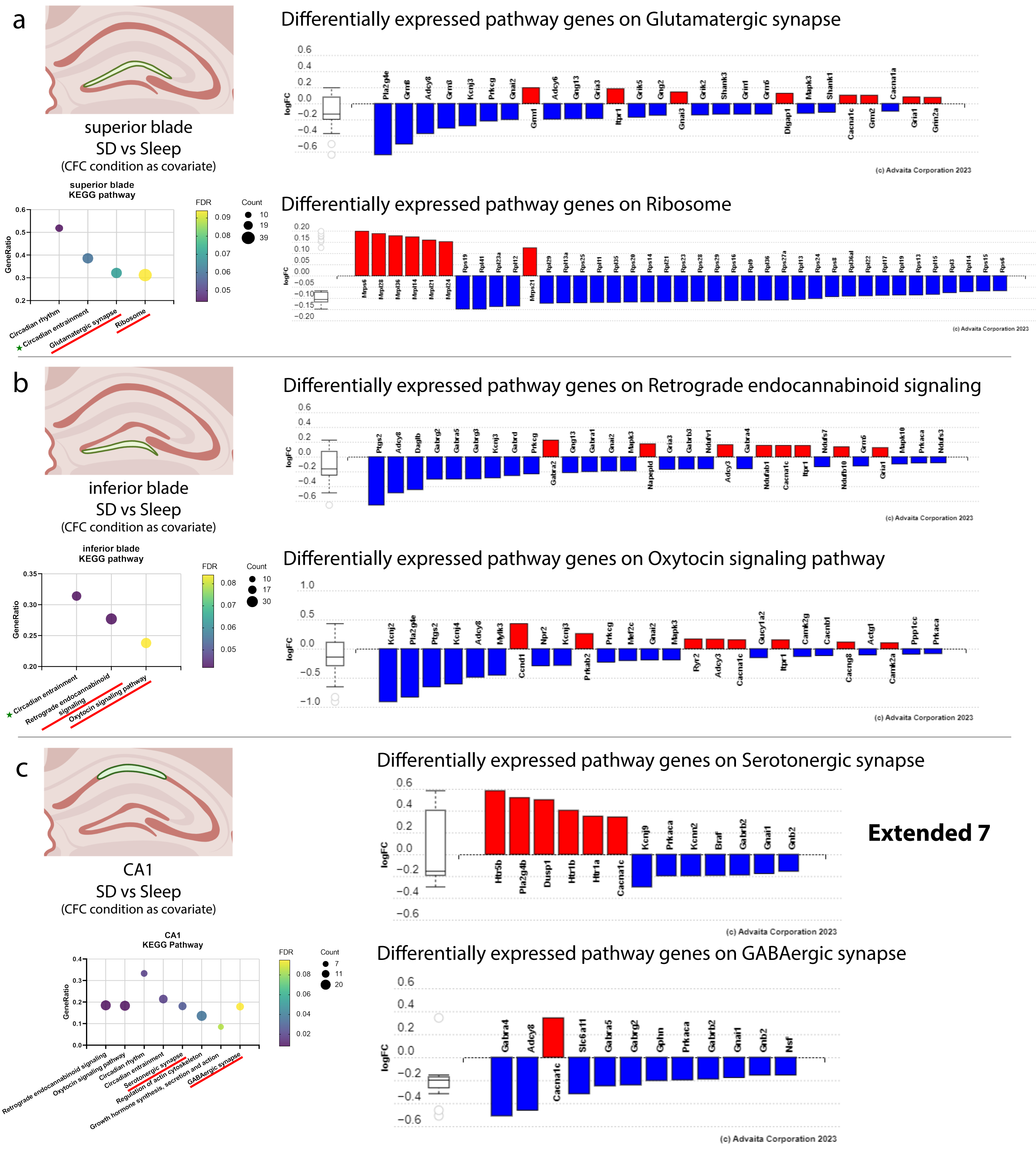
a
superior blade
SD vs Sleep
(CFC condition as covariate)
superior blade
KEGG pathway
FDR
Count
GeneRatio
Circadian rhythm
Circadian entrainment
Glutamatergic synapse
Ribosome
Differentially expressed pathway genes on Glutamatergic synapse
logFC
(c) Advaita Corporation 2023
Differentially expressed pathway genes on Ribosome
b
inferior blade
SD vs Sleep
(CFC condition as covariate)
inferior blade
KEGG pathway
Circadian entrainment
Retrograde endocannabinoid signaling
Oxytocin signaling pathway
Differentially expressed pathway genes on Retrograde endocannabinoid signaling
Differentially expressed pathway genes on Oxytocin signaling pathway
c
CA1
SD vs Sleep
(CFC condition as covariate)
CA1
KEGG Pathway
Retrograde endocannabinoid signaling
Oxytocin signaling pathway
Circadian rhythm
Circadian entrainment
Serotonergic synapse
Regulation of actin cytoskeleton
Growth hormone synthesis, secretion and action
GABAergic synapse
Differentially expressed pathway genes on Serotonergic synapse
Extended 7
Differentially expressed pathway genes on GABAergic synapse

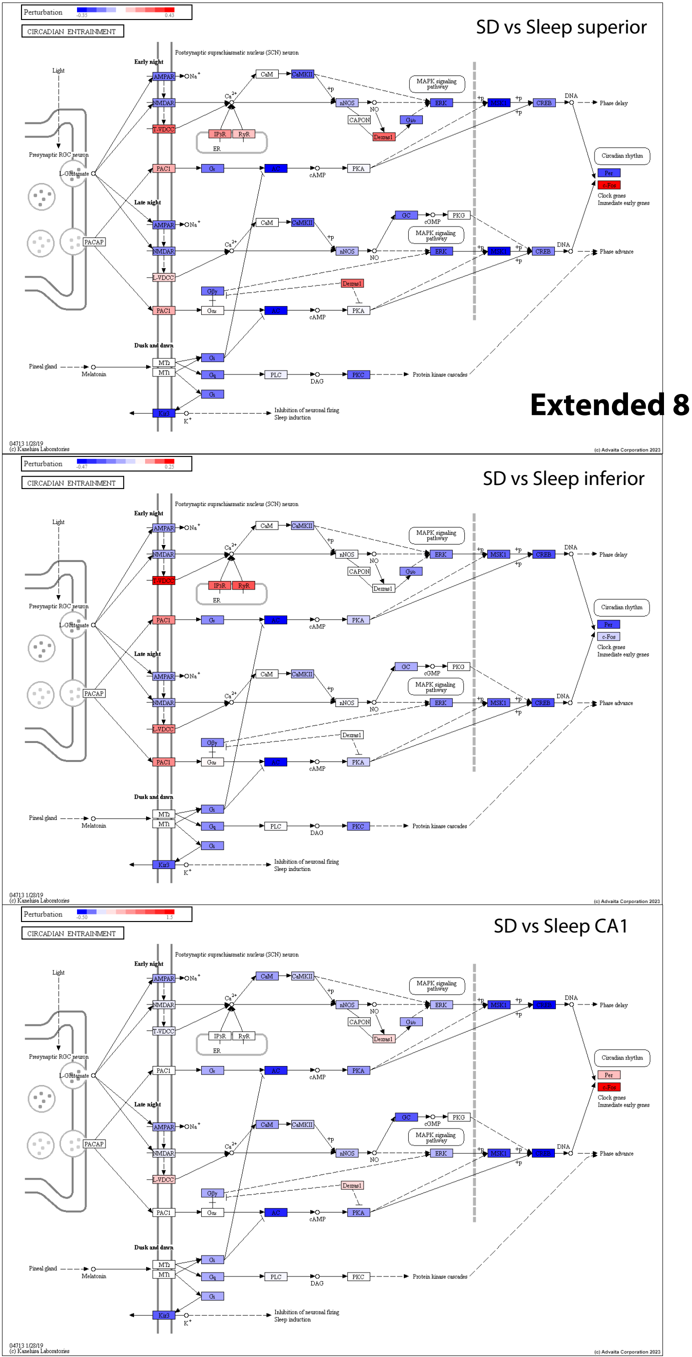
Extended 8
SD vs Sleep superior
SD vs Sleep inferior
SD vs Sleep CA1
Perturbation
-0.35
0.43
-0.47
0.25
-0.50
1.5
CIRCADIAN ENTRAINMENT
Postsynaptic suprachiasmatic nucleus (SCN) neuron
Early night
Late night
Dusk and dawn
Light
Presynaptic RGC neuron
L-Glutamate
PACAP
Pineal gland
Melatonin
AMPAR
NMDAR
T-VDCC
L-VDCC
PAC1
MT2
MT1
Kir3
CaM
CaMKII
nNOS
NO
CAPON
Dexras1
Gi/o
IP3R
RyR
ER
Gs
Gi
Gq
Gαs
Gβγ
AC
cAMP
PKA
PLC
DAG
PKC
GC
cGMP
PKG
ERK
MSK1
CREB
DNA
MAPK signaling pathway
Phase delay
Phase advance
Circadian rhythm
Per
c-Fos
Clock genes
Immediate early genes
Protein kinase cascades
Inhibition of neuronal firing
Sleep induction
04713 1/28/19
(c) Kanehisa Laboratories
(c) Advaita Corporation 2023

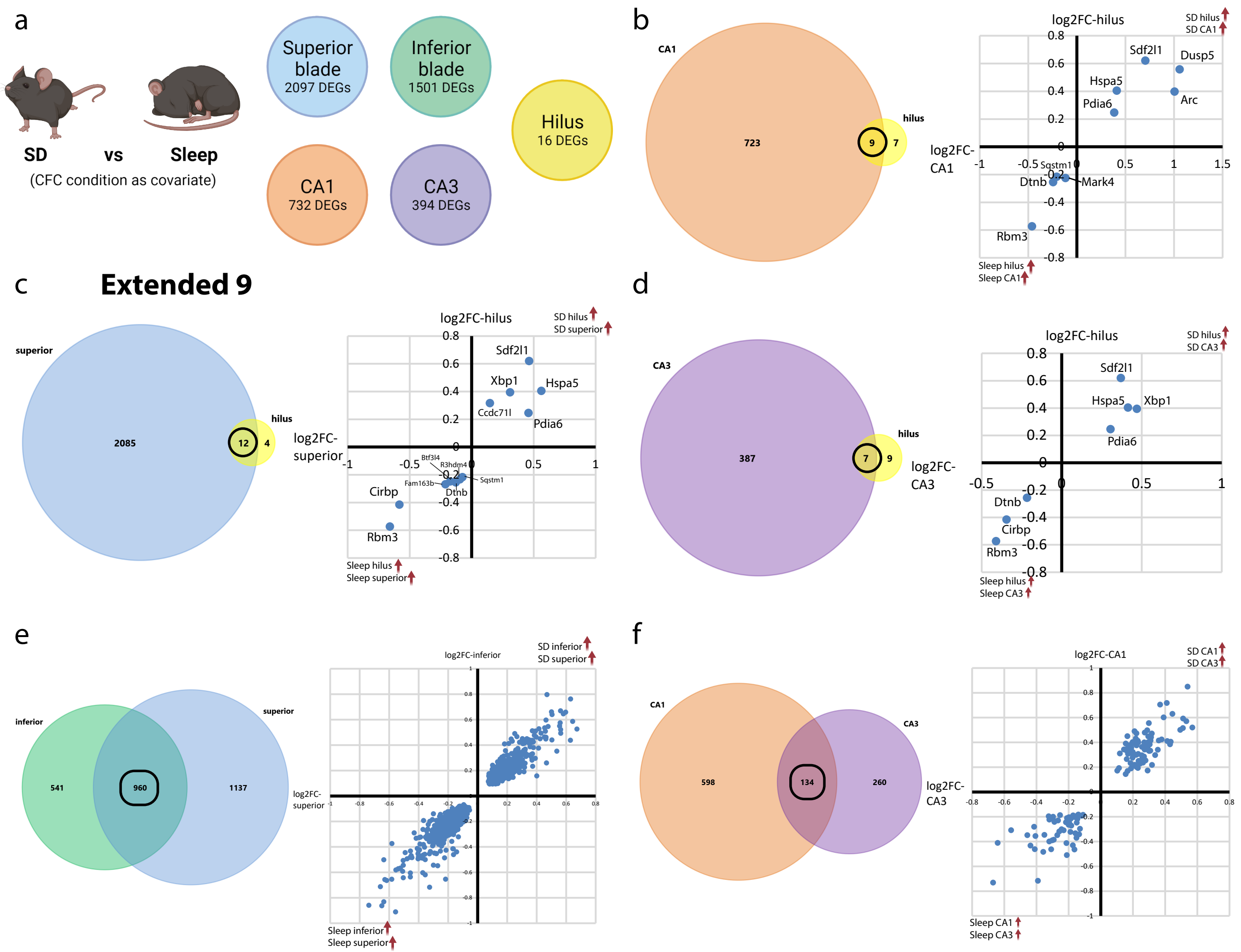
a
SD
vs
Sleep
(CFC condition as covariate)
Superior blade
2097 DEGs
Inferior blade
1501 DEGs
Hilus
16 DEGs
CA1
732 DEGs
CA3
394 DEGs
b
CA1
723
9
7
hilus
log2FC-hilus
SD hilus
SD CA1
log2FC-CA1
Sleep hilus
Sleep CA1
Sdf2l1
Dusp5
Hspa5
Arc
Pdia6
Sqstm1
Dtnb
Mark4
Rbm3
c
Extended 9
superior
2085
12
4
hilus
log2FC-hilus
SD hilus
SD superior
log2FC-superior
Sleep hilus
Sleep superior
Sdf2l1
Xbp1
Hspa5
Ccdc71l
Pdia6
Btf3l4
R3hdm4
Fam163b
Sqstm1
Dtnb
Cirbp
Rbm3
d
CA3
387
7
9
hilus
log2FC-hilus
SD hilus
SD CA3
log2FC-CA3
Sleep hilus
Sleep CA3
Sdf2l1
Hspa5
Xbp1
Pdia6
Dtnb
Cirbp
Rbm3
e
inferior
superior
541
960
1137
log2FC-inferior
SD inferior
SD superior
log2FC-superior
Sleep inferior
Sleep superior
f
CA1
CA3
598
134
260
log2FC-CA1
SD CA1
SD CA3
log2FC-CA3
Sleep CA1
Sleep CA3

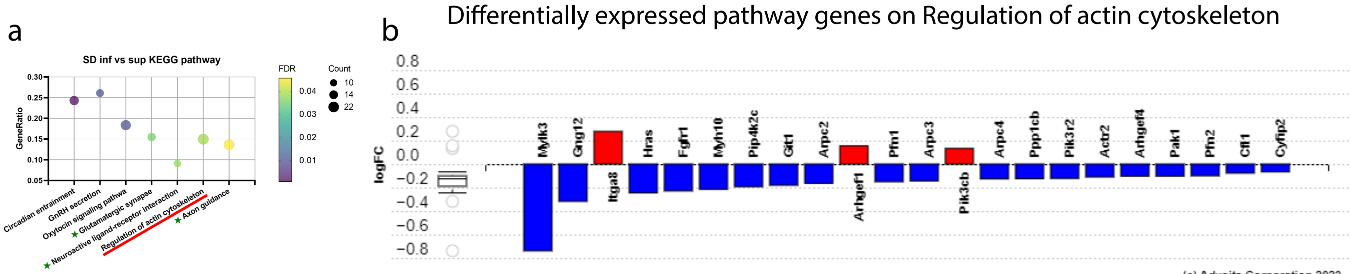
a
SD inf vs sup KEGG pathway
GeneRatio
0.30
0.25
0.20
0.15
0.10
0.05
Circadian entrainment
GnRH secretion
Oxytocin signaling pathwa
Glutamatergic synapse
Neuroactive ligand-receptor interaction
Regulation of actin cytoskeleton
Axon guidance
FDR
0.04
0.03
0.02
0.01
Count
10
14
22
b
Differentially expressed pathway genes on Regulation of actin cytoskeleton
logFC
0.8
0.6
0.4
0.2
0.0
−0.2
−0.4
−0.6
−0.8
Mylk3
Gng12
Itga8
Hras
Fgfr1
Myh10
Pip4k2c
Git1
Arpc2
Arhgef1
Pfn1
Arpc3
Pik3cb
Arpc4
Ppp1cb
Pik3r2
Actr2
Arhgef4
Pak1
Pfn2
Cfl1
Cyfip2
(c) Advaita Corporation 2023

**Extended 10**

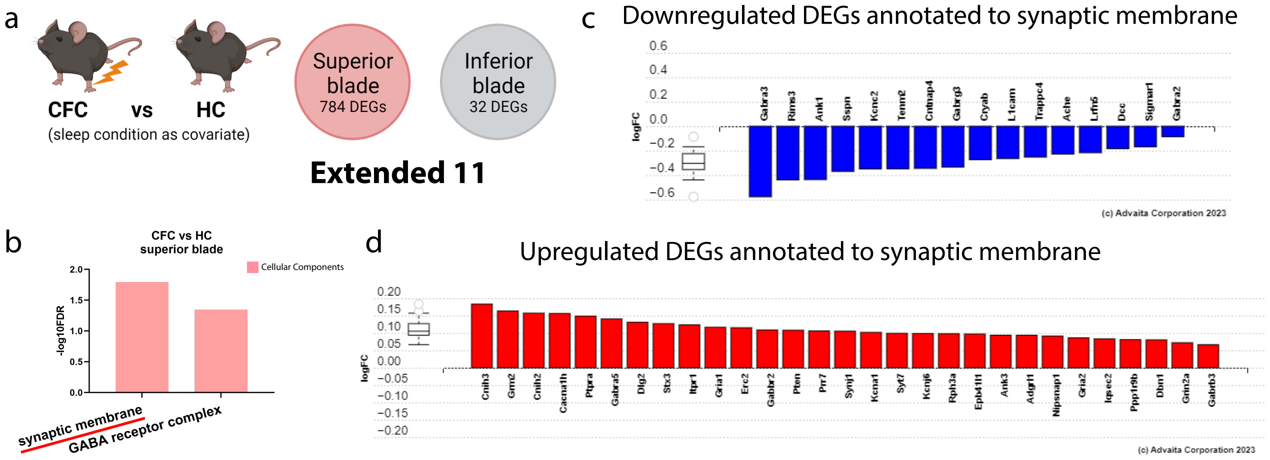
a
CFC vs HC
(sleep condition as covariate)
Superior blade
784 DEGs
Inferior blade
32 DEGs
Extended 11
b
CFC vs HC
superior blade
Cellular Components
-log10FDR
2.0
1.5
1.0
0.5
0.0
synaptic membrane
GABA receptor complex
c
Downregulated DEGs annotated to synaptic membrane
logFC
0.6
0.4
0.2
0.0
−0.2
−0.4
−0.6
Gabra3
Rims3
Ank1
Sspn
Kcnc2
Tenm2
Cntnap4
Gabrg3
Cryab
L1cam
Trappc4
Ache
Lrfn5
Dcc
Sigmar1
Gabra2
(c) Advaita Corporation 2023
d
Upregulated DEGs annotated to synaptic membrane
logFC
0.20
0.15
0.10
0.05
0.00
−0.05
−0.10
−0.15
−0.20
Cnih3
Grm2
Cnih2
Cacna1h
Ptpra
Gabra5
Dlg2
Stx3
Itpr1
Gria1
Erc2
Gabbr2
Pten
Prr7
Synj1
Kcna1
Syt7
Kcnj6
Rph3a
Epb41l1
Ank3
Adgrl1
Nipsnap1
Gria2
Iqsec2
Ppp1r9b
Dbn1
Grin2a
Gabrb3
(c) Advaita Corporation 2023

### Gaine et al., 2021 (hippocampal transcripts)

a

### Lyons et al., 2020 (hippocampal Camk2a transcripts)

b

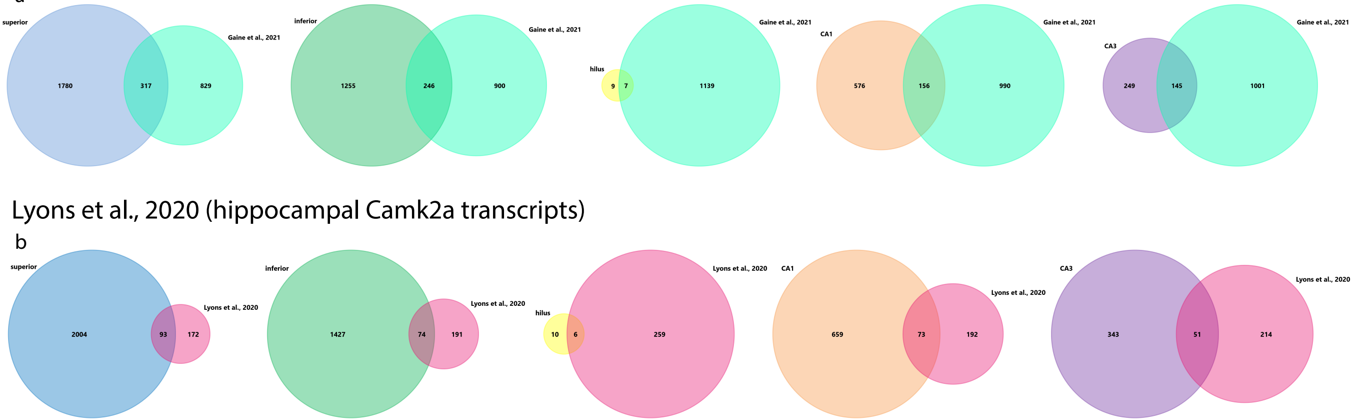

### Delorme et al., 2021 (Hippocampal input, Camk2a, and pS6 transcripts altered by SD)

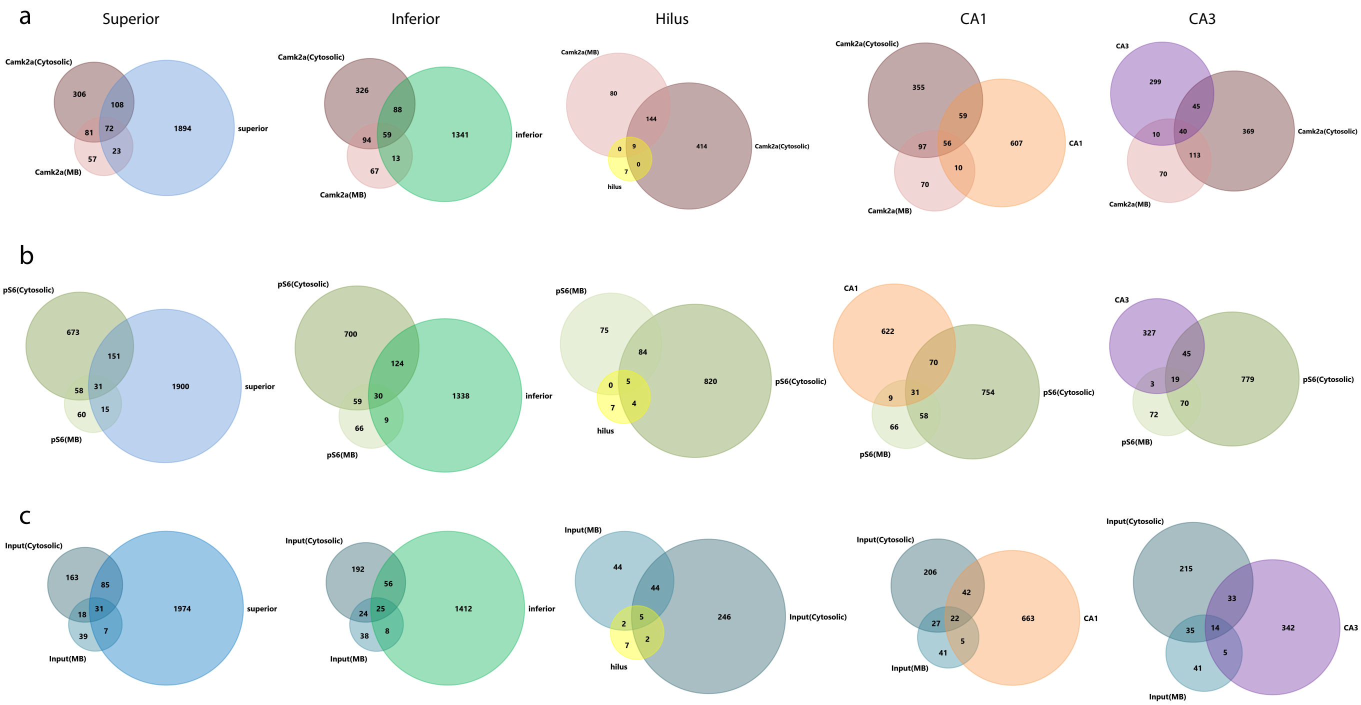
